# Direct CDK1 engagement underlies carbaryl-induced genotoxic cell cycle arrest

**DOI:** 10.64898/2026.10.01.756001

**Authors:** Ravikant Piyush, Sumedh Tayade, Swapnil Rohidas Shinde

## Abstract

Carbaryl, a widely used carbamate pesticide, has been epidemiologically associated with genotoxic and carcinogenic outcomes in exposed human populations, including elevated cancer risk at several tissue sites and, in some agricultural communities, chronic tubulointerstitial kidney disease of unknown etiology (CKDu). However, the cellular mechanisms underlying these associations, and whether carbaryl acts uniformly across tissues, remain poorly defined. Screening a panel of cell lines of distinct species and lineage, we found that carbaryl elicits pronounced, dose-dependent cytotoxicity and cell cycle arrest selectively in a renal tubular epithelial line (IMCD3) and, to a lesser extent, a cervical epithelial line (HeLa), while a fibroblast line (NIH3T3) was unaffected. In IMCD3 cells, carbaryl induces S- and G2/M-phase arrest through direct binding to DNA and consequent DNA damage, triggers *Bcl-2*-mediated late apoptosis independently of reactive oxygen species, and drives irreversible mitotic arrest. Molecular dynamics simulations, cellular thermal shift assays, and kinase measurements converge on CDK1 as a direct target of carbaryl: carbaryl destabilises the CDK1 active site and inhibits its catalytic activity, whereas CDK2 is engaged through an opposing, rigidifying mode with no functional consequence. This asymmetry provides a coherent mechanistic basis for the combined S-phase and G2/M arrest observed. Together, these findings establish carbaryl as a genotoxic, cell-cycle-disrupting agent with tissue-selective potency acting directly on CDK1, and provide a cell-based mechanistic framework for interpreting its reported associations with renal and other tissue pathologies in exposed human populations.

## Introduction

Environmental and occupational exposure to agrochemicals is an increasingly recognised contributor to human disease, and understanding how individual compounds compromise cellular integrity is essential for assessing these risks. Since the fidelity of cell division underlies healthy tissue maintenance, xenobiotics that perturb cell cycle progression can translate into organ-level pathology ranging from carcinogenesis to progressive degenerative disease.

Multiple checkpoints monitor the cell cycle for DNA damage, replication stress, and spindle or chromosome-segregation defects [1]; when severely compromised, cells are eliminated via mitotic catastrophe, a tumour-suppressive process that gates entry into apoptosis, necrosis, autophagic death, or senescence depending on cellular context [2–4]. This regulation is enforced by cyclin-CDK complexes acting in a phase-specific manner: cyclin D-CDK4/6 drives G1 entry, cyclin E/A-CDK2 drives S phase, and cyclin B-CDK1 drives mitosis [5,6], with CDK1 and CDK2, despite sharing over 60% sequence identity, performing largely non-redundant roles in somatic cell division. CDK2 mainly governs the G1/S transition, CDK1 governs mitosis, and APC/C-mediated cyclin degradation resets activity to zero in daughter cells [1].

Carbaryl (1-naphthol N-methylcarbamate) is among the most widely used carbamate insecticides globally, with millions of pounds applied annually in the United States alone [7,8], though its use has been restricted or banned in the European Union and India [9,10]. Beyond its well-characterised cholinesterase-inhibiting neurotoxicity, carbaryl exposure has been linked to oxidative stress, endocrine disruption via estrogen receptor activation, and genotoxicity, including through formation of carcinogenic N-nitroso derivatives **(Supplementary information; Table S1)** [11]. Epidemiological studies report elevated cancer risk in carbaryl-exposed populations, consistently for melanoma and gliomas but less consistently for non-Hodgkin lymphoma [12], alongside reproductive toxicity and elevated chromosomal aberrations and micronuclei frequency in biomonitored agricultural workers, indicating that carbaryl’s genotoxic potential manifests across multiple tissue systems in humans.

Despite this evidence, the mechanistic basis by which carbaryl perturbs cell cycle progression, and whether this effect is uniform across tissues, remains poorly defined. Because xenobiotic metabolism via cytochrome P450 (CYP) enzymes varies considerably across cell types [13,14], carbaryl’s cytotoxic and cell-cycle-disrupting effects may depend on the CYP expression profile of the exposed tissue, a question not yet systematically addressed using a comparative cell-based model. The kidney is one compelling candidate tissue in this context: as the principal organ responsible for xenobiotic filtration and excretion, renal tubular epithelium is directly exposed to carbaryl at high local concentrations, and agrochemical exposure has been implicated in chronic tubulointerstitial kidney disease of unknown etiology (CKDu), an emerging global health burden affecting agricultural communities across South Asia, Central America, and parts of Africa, including well-documented endemic clusters such as Uddanam nephropathy in Andhra Pradesh, India [15–17]. Given that CKDu pathology is centered on tubular epithelial injury rather than glomerular damage, renal tubular epithelial cell models are well suited to mechanistically probe whether carbaryl can directly compromise tubular cell integrity and cell cycle fidelity, offering a cell-based correlate to these epidemiological associations.

Here, we investigated the effect of carbaryl on cell cycle progression across three cell lines of distinct species and tissue lineage (IMCD3, NIH3T3, HeLa) to determine whether carbaryl elicits lineage-dependent effects correlating with CYP expression. Having identified IMCD3 as the most sensitive line, we dissected the underlying mechanism, examining direct carbaryl-DNA interaction and damage, ROS-independent *Bcl-2*-mediated apoptosis, irreversibility of mitotic arrest, and direct modulation of CDK1 kinase activity governing the S and G2/M checkpoints.

## Results

### Carbaryl exhibits lineage-selective cytotoxicity, cell cycle arrest, and promotes late apoptosis

To identify which cell types are most susceptible to carbaryl, we screened a panel of cell lines (IMCD3, NIH3T3, HeLa) of distinct origin. Viability and cytotoxicity assays revealed a dose-dependent response only in IMCD3 and HeLa cells, with EC50/IC50 values of 23.4/24 µM and 112/129 µM, respectively; NIH3T3 showed no response **(Figure 1A; S1A** and **B)**.

**Figure 1.**
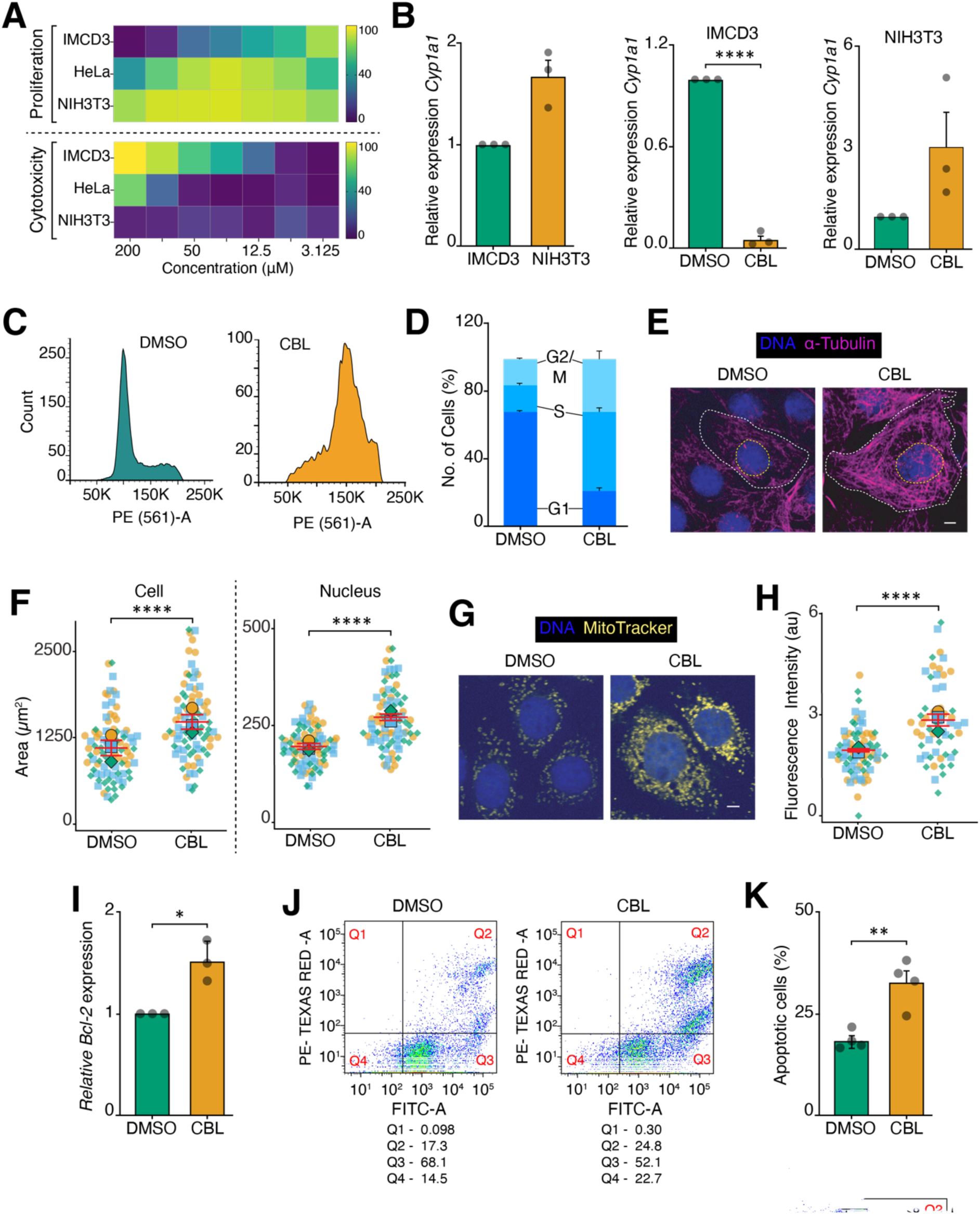
Carbaryl exposure induces cell cycle arrest, cellular hypertrophy, and late apoptosis in a lineage-dependent manner. **(A)** Heatmaps showing dose-dependent effects of carbaryl on proliferation (top) and cytotoxicity (bottom) in IMCD3, HeLa, and NIH3T3 cells across a concentration range (3.125 - 200 µM), illustrating differential sensitivity across the three cell lines. **(B)** qRT-PCR analysis of *Cyp1a1* in IMCD3 and NIH3T3 cells treated with DMSO or carbaryl. Left: Gene expression in IMCD3 was set to 1. Middle: Gene expression in the DMSO-treated condition was set to 1. Right: Gene expression in the DMSO-treated condition was set to 1. Data represent the mean ± SEM from three independent experiments. **(C)** Representative flow cytometry histograms of DNA content in IMCD3 cells treated with DMSO or carbaryl; a shift in cell cycle distribution upon carbaryl treatment was observed with S/G2-M accumulation. **(D)** Quantification of cell cycle phase distribution (G1, S, G2/M) in DMSO- versus carbaryl-treated IMCD3 cells. Data represent mean ± SEM from three independent experiments. **(E)** Representative immunofluorescence images of IMCD3 cells stained for DNA (blue) and α-tubulin (magenta) following DMSO or carbaryl treatment. Dashed white lines outline cell boundaries; dashed yellow lines outline nuclei. Scale bar: 5 µm. **(F)** Quantification of cell area (left) and nuclear area (right) in DMSO- versus carbaryl-treated IMCD3 cells. In the superplots, three independent experimental data points are represented by three different shapes, with colours indicated. Each square, circle, or triangle represents a data point, and the larger version of its shape represents the mean. Data represent all data points from three independent experiments, mean ±SEM. DMSO n = 89, carbaryl n = 84. **(G)** Representative fluorescence images of DMSO- and CBL-treated IMCD3 cells stained with MitoTracker (yellow) and DNA (blue). Scale bar: 5 µm. **(H)** Quantification of mitochondrial membrane intensity (MitoTracker fluorescence) in DMSO- versus carbaryl-treated IMCD3 cells. Superplot convention as in F. Data represent all data points from three independent experiments, mean ±SEM. DMSO n = 75, carbaryl n = 55. **(I)** qRT-PCR analysis of *Bcl-2* in IMCD3 cells treated with DMSO or carbaryl. Gene expression in the DMSO-treated condition was set to 1. Data represent the mean ± SEM from three independent experiments. **(J)** Representative flow cytometry dot plots of Annexin V-FITC/PI staining in DMSO- and carbaryl-treated IMCD3 cells. **(K)** Quantification of the apoptotic cell population (%) from Annexin V/PI flow cytometry in DMSO- versus carbaryl-treated IMCD3 cells. Data represent mean ± SEM from four independent experiments. Fluorescence intensities are expressed in arbitrary units (a.u.). Asterisks denote statistical significance as determined by unpaired Student’s t-test: **** p ≤ 0.0001; *** p ≤ 0.001; ** p ≤ 0.01; * p ≤ 0.05.

Carbaryl is metabolised predominantly via CYP-mediated hydroxylation [18,19]; we hypothesised that differential CYP expression underlies this divergent sensitivity. qRT-PCR showed that basal *Cyp1a1* and *Cyp1a2* expression was significantly lower in IMCD3 than in NIH3T3, and that carbaryl treatment further downregulated *Cyp1a1* in IMCD3 while upregulating it in NIH3T3, with the opposite trend for *Cyp1a2* in each line **(Figure 1B** and **S1C)**. These findings demonstrate that carbaryl exposure differentially regulates CYP expression across cell lines.

Next, we assessed the effect of carbaryl on cell cycle progression to determine whether cell proliferation is inhibited by cell cycle arrest. Flow cytometric cell cycle analysis after 48 h of treatment showed that carbaryl induces S- and G2/M-phase arrest in IMCD3 and HeLa cells. In IMCD3, S-phase and G2/M populations increased from 15.6 ± 2.1% and 15.9 ± 1.0% (control) to 46.9 ± 4.6% and 31.5 ± 8.1% (carbaryl), respectively **(Figure 1C** and **D)**. In HeLa, they increased from 13.6 ± 4.3% and 3.6 ± 1.9% to 32.5 ± 4.2% and 13.9 ± 1.8%, respectively **(Figure S1D)**.

Interestingly, carbaryl-treated IMCD3 cells showed a marked, proportionate increase in both cell and nuclear area, with an unchanged nucleus-to-cell ratio **(Figure 1E-F** and **S1E)**, consistent with a cellular hypertrophy phenotype typical of cell cycle arrest rather than selective nuclear swelling. Given the established link between xenobiotic-induced cell cycle perturbation and mitochondrial stress, we assessed mitochondrial membrane potential using MitoTracker Red staining. Carbaryl significantly hyperpolarised mitochondrial membrane potential (MitoTracker), but not intracellular ROS (unlike the doxorubicin positive control), indicating that this mitochondrial change is ROS-independent **(Figure 1G-H** and **S1F)**.

Moreover, *Bcl-2* expression was significantly increased in carbaryl-treated cells, consistent with cells committing to an apoptotic fate **(Figure 1I)**. Annexin V/PI flow cytometry confirmed a significant increase in the overall apoptotic population, with selective enrichment of the late-apoptotic fraction, indicating that carbaryl-treated cells progress well along the apoptotic cascade rather than merely initiating early apoptotic signalling **(Figure 1J-K)**.

### DNA damage-induced cell cycle arrest upon carbaryl exposure

To identify the cause of cell cycle arrest upon carbaryl exposure, we assessed whether carbaryl induces DNA damage leading to cell cycle block. We checked the nuclear accumulation of phosphorylated γH2AX (Ser-139), a well-established marker for DNA damage. Phospho-γH2AX was significantly increased in IMCD3 but unchanged in HeLa and NIH3T3 **(Figure 2A-B and S2A-D)**. This was confirmed by immunoblotting in IMCD3 cells, corroborating the immunostaining result **(Figure 2C)**.

**Figure 2.**
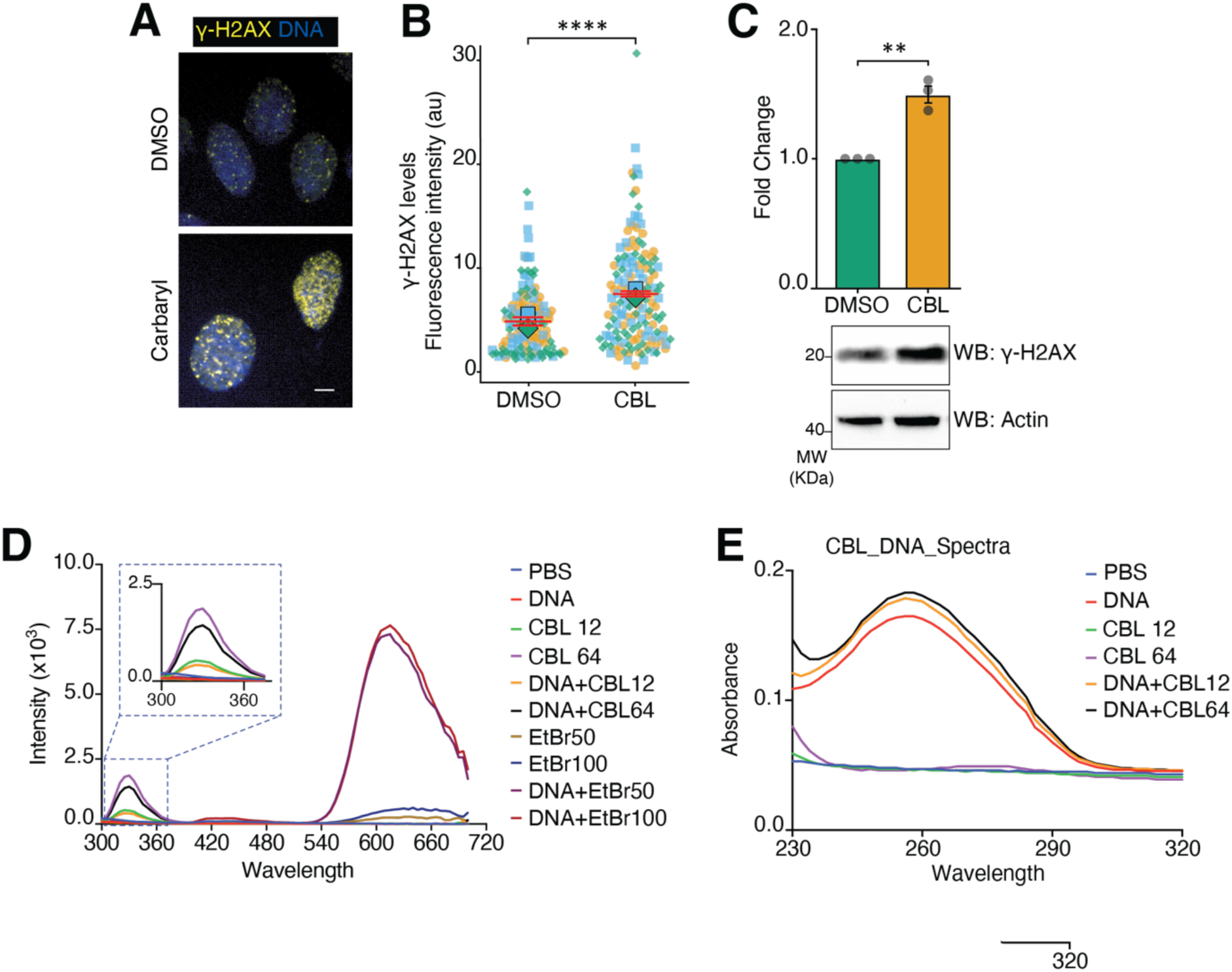
Carbaryl induces DNA damage via direct interaction. **(A)** Representative immunofluorescence images of IMCD3 cells stained for γ-H2AX (yellow) and DNA (blue) following DMSO or carbaryl treatment. Scale bar: 5 µm. **(B)** Quantification of γ-H2AX fluorescence intensity per nucleus in DMSO- versus carbaryl-treated IMCD3 cells. In the superplots, three independent experimental data points are represented by three different shapes, with colours indicated. Each square, circle, or triangle represents a data point, and the larger version of its shape represents the mean. Data represent all data points from three independent experiments, mean ±SEM. DMSO n = 136, carbaryl n = 151. **(C)** IMCD3 cells were lysed, and the lysates were probed for γ-H2AX and actin (loading control) via immunoblotting. γ-H2AX signal intensities were measured and normalised against the loading control. γ-H2AX expression in the DMSO condition was set to 1. The data represent the mean ±SEM of three independent experiments. **(D)** Fluorescence emission spectra scan (300–720 nm) of carbaryl (12 and 64 µM) with DNA (100 ng). Ethidium bromide (50 and 100 nM) was used as a positive control. Varying concentrations of carbaryl were incubated with DNA. The inset shows a magnified view of the carbaryl emission peak region (300–360 nm). **(E)** UV-Vis absorbance spectra (230–320 nm) of DNA (100 ng) incubated with different concentrations of carbaryl (12 and 64 µM). Fluorescence intensities are expressed in arbitrary units (a.u.). Asterisks denote statistical significance as determined by unpaired Student’s t-test: **** p ≤ 0.0001; *** p ≤ 0.001; ** p ≤ 0.01; * p ≤ 0.05.

Although previous studies indicated that carbaryl induces DNA damage via reactive oxygen species (ROS) generation and direct DNA binding, we evaluated whether these mechanisms were active at our specific concentration. Fluorescence spectroscopy with DCFHDA dye revealed no significant ROS production **(Figure S1F)**. However, UV-vis and fluorescence spectroscopy demonstrated shifts in the peak intensities of DNA and carbaryl, respectively, confirming a direct binding interaction **(Figure 2D-E)**. Consequently, this direct binding interaction represents the primary mechanism underlying the observed DNA damage independent of ROS.

### Carbaryl induces severe mitotic arrest

Given persistent DNA damage, we asked whether the G2/M-gated population identified by flow cytometry reflected cells held in G2 or stalled within mitosis itself, a distinction flow cytometry cannot resolve on DNA content alone. To investigate carbaryl’s impact on the microtubule network, we measured total alpha- and gamma-tubulin levels and found no change in microtubule levels in IMCD3 cells **(Figure S3A-B)**. We further confirmed this by staining phospho-Histone H3, a mitosis marker, showing an increased mitotic population in carbaryl-treated IMCD3 cells after 48 hours, with no change in HeLa and NIH3T3 cells **(Figure 3A-B** and **S3C-F)**, indicating that a substantial fraction of the IMCD3 G2/M population is arrested specifically within mitosis rather than in G2. This arrest was irreversible: a significant mitotic population persisted in IMCD3 even 16 h after carbaryl withdrawal **(Figure 3C-E; S4** and **S5)**. A microtubule cold-shock recovery assay showed unaltered polymerisation/depolymerisation dynamics, indicating that carbaryl does not act through the microtubule network or the microtubule-organising centre to produce this arrest **(Figure S6)**.

**Figure 3.**
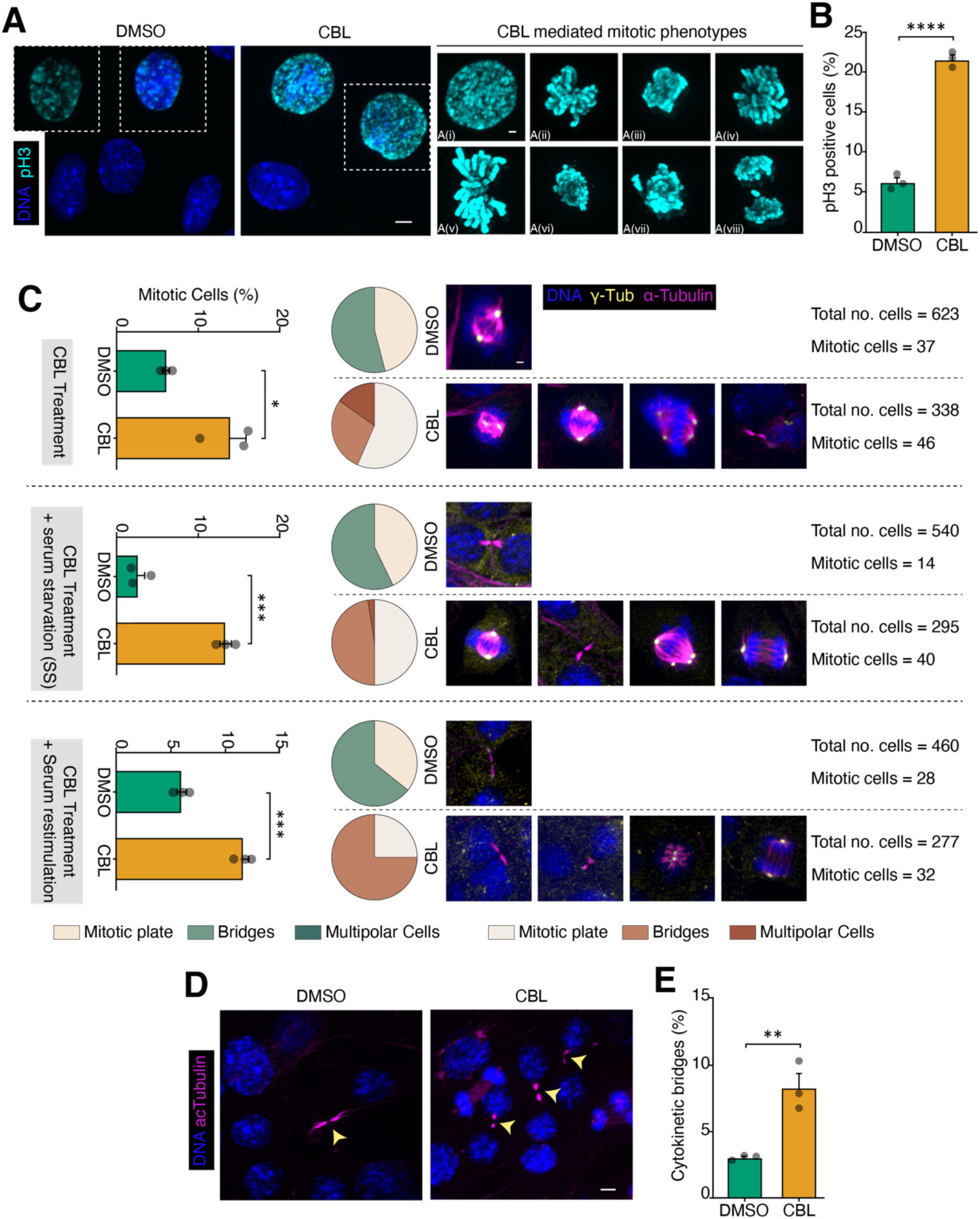
Carbaryl induces irreversible mitotic arrest. **(A)** Representative immunofluorescence images of IMCD3 cells stained for phospho-histone H3 (pH3, cyan) and DNA (blue) following DMSO or carbaryl treatment, illustrating the range of carbaryl-induced mitotic chromosome phenotypes (panels B(i)-B(viii)). Scale bar: 5 µm. **(B)** Quantification of the percentage of pH3-positive (mitotic) cells in DMSO- versus carbaryl-treated IMCD3 cells. Data represent the mean ± SEM from three independent experiments. **(C)** Quantification and classification of mitotic defects in IMCD3 cells under three experimental conditions: 48 hours of carbaryl treatment (top), 48 hours of carbaryl treatment followed by 16 hours of serum starvation (+SS, middle), and 48 hours of carbaryl treatment followed by 16 hours of serum re-stimulation (+Serum, bottom). *Left:* Bar graphs showing the percentage of mitotic cells in DMSO- versus carbaryl-treated populations under each condition. Data represent mean ± SEM from three independent experiments. *Middle:* Pie charts showing the proportional distribution of mitotic phenotypes, i.e., mitotic plate (normal metaphase alignment), bridges (chromatin/anaphase bridges), and multipolar cells (aberrant multipolar spindles) in DMSO- versus carbaryl-treated cells for each condition. *Right:* Representative immunofluorescence images of mitotic IMCD3 cells stained for DNA (blue), γ-tubulin (yellow), and α-tubulin (magenta) in DMSO- and carbaryl-treated cells for each condition. **(D)** Representative immunofluorescence images of IMCD3 cells stained for DNA (blue) and αcTubulin (magenta) following DMSO or carbaryl treatment for 48 hours followed by 16 hours of serum starvation. Yellow arrowheads indicate cytokinetic bridges identified within each field. Scale bar: 5 µm. **(E)** Quantification of the percentage of cytokinetic bridges in DMSO- versus carbaryl-treated IMCD3 cells. Data represent the mean ± SEM from three independent experiments. Asterisks denote statistical significance as determined by unpaired Student’s t-test: **** p ≤ 0.0001; *** p ≤ 0.001; ** p ≤ 0.01; * p ≤ 0.05.

### Carbaryl modulates a key regulator of the cell cycle: CDK1

Given the severity of cell cycle arrest, we hypothesised that carbaryl directly engages core cell cycle regulators. Using molecular docking followed by 500 ns molecular dynamics (MD) simulations, we tested carbaryl binding to the key regulators CDK1, CDK2, and CDK4 during cell cycle progression.

Carbaryl showed binding to CDK1 and CDK2 **(Figure S7A-B)**. In both complexes, carbaryl formed stabilising hydrophobic and π-π interactions within the active site, with no change in binding-site surface potential relative to the apo form **(Figure 4A**, **5A, S7C** and **S8B)**. However, the two kinases responded oppositely at the structural level. Carbaryl binding shifted CDK1 from a single, well-defined RMSD-Rg basin to a higher-RMSD, higher-Rg, bimodal ensemble **(Figure 4B)**, with increased flexibility across several loop clusters (including regions near residues 18-22, 87-89, 140-150, and 230-250) **(Figure 4C-E)** and increased surface exposure near residue 40, plus secondary sites near residues 170-180 and the C-terminus **(Figure 4F and S7E-F)**, consistent with local loop opening rather than compaction; the free-energy landscape confirmed a shift of the global minimum toward this higher-Rg state **(Figure 4G and S7G)**. CDK2 showed the opposite trend: carbaryl binding shifted the ensemble to a tighter, lower-Rg basin, non-overlapping with the apo state **(Figure 5B, G)**, with net rigidification (reduced flexibility across residues 50-100 and near residue 300, alongside one localised region of increased mobility near residues 150-190) **(Figure 5C-E)** and a dominant burial event near residue 20, indicating ligand-induced stabilisation of a more compact conformation despite this local flexibility **(Figure 5F)**. For both kinases, global conformational entropy was preserved despite these local perturbations **(Figure S7H and S8G)**.

**Figure 4.**
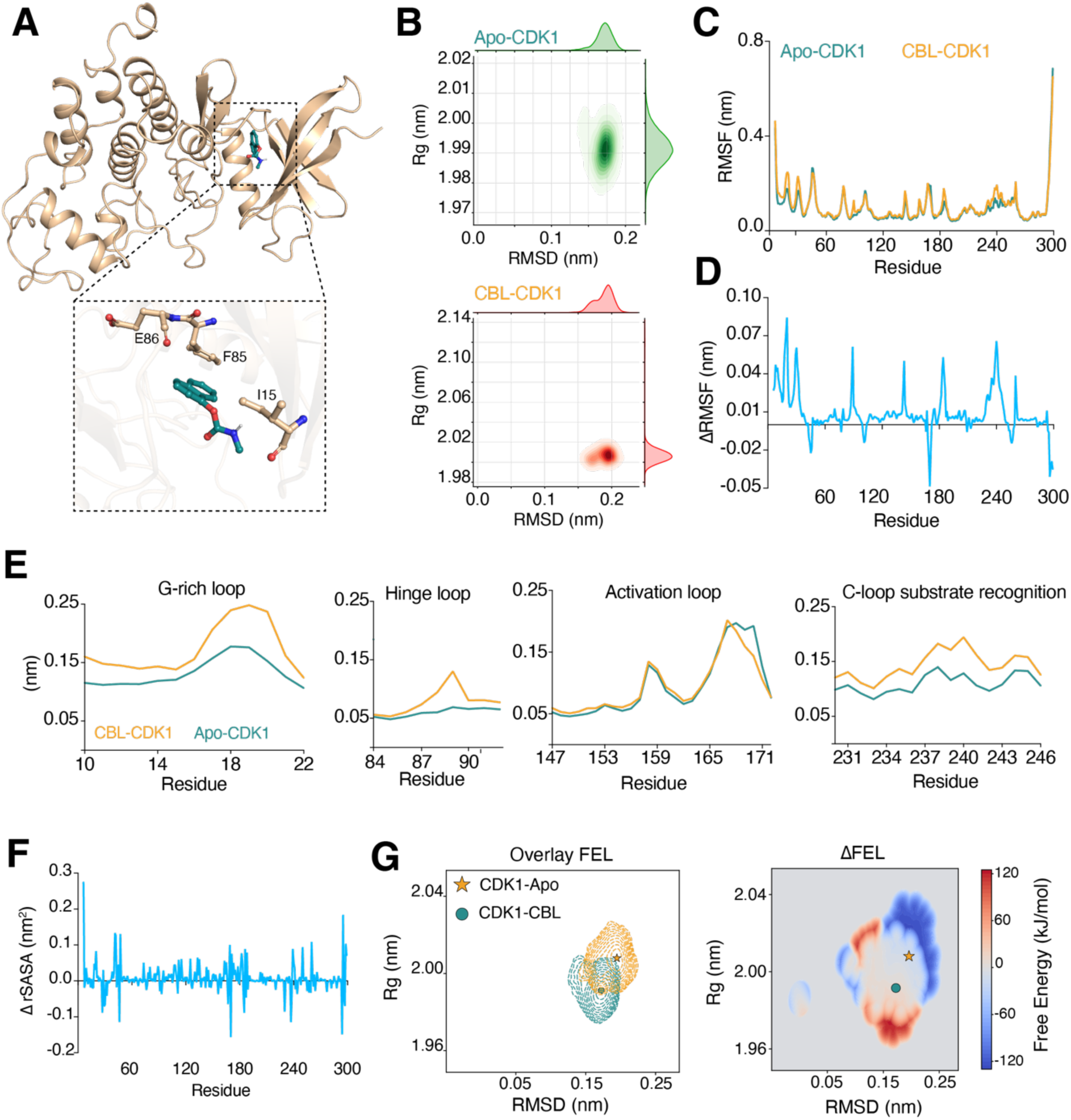
Carbaryl stably interacts with CDK1 and induces conformational changes. **(A)** Representative structure of the CDK1: carbaryl complex after a 500 ns MD simulation. The inset shows key residues of CDK1 involved in interactions with carbaryl throughout the 500 ns simulation. **(B)** RMSD versus radius of gyration (Rg) plot of CDK1 in the apo (green) and carbaryl-bound (red) forms, demonstrating the structural stability and compactness of CDK1 throughout the 500 ns MD simulation. **(C)** Root mean square fluctuation (RMSF) of CDK1 residues in the apo (green) and carbaryl-bound (wheat) forms. **(D)** Delta RMSF (ΔRMSF) for individual CDK1 residues (nm), illustrating changes in residue fluctuations. **(E)** The zoomed view of the RMSF of the G-rich loop (residues 10-22), hinge loop (residues 78-92), activation loop (residues 147-171), and C-loop substrate recognition (residues 231-246) of CDK1 in the apo (green) and carbaryl-bound (wheat) forms. **(F)** Delta relative solvent-accessible surface area (ΔrSASA) expressed as a percentage for individual CDK1 residues, illustrating changes in solvent accessibility. **(G)** Two-dimensional overlay and change in free energy landscape (ΔFEL) of CDK1 in the apo and carbaryl-bound forms. The global minimum energy state in the apo form is indicated by a filled circle (●) and in the carbaryl-bound form by a filled star (★). The MD simulations and the corresponding analyses were performed in triplicate, and the representative graphs show the mean across three independent simulations for each parameter analysed.

**Figure 5.**
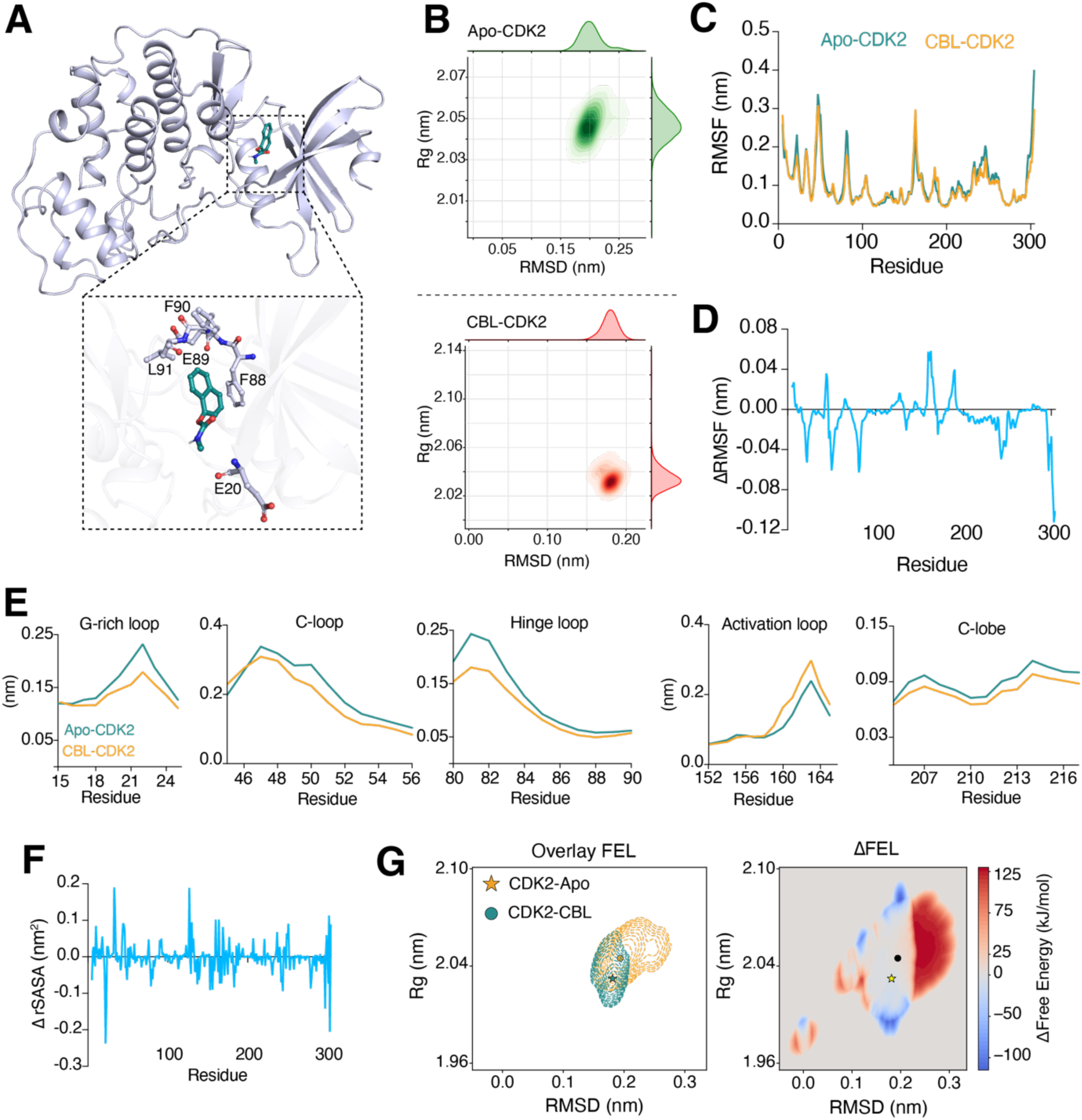
Carbaryl stably interacts with CDK2 and induces conformational changes. **(A)** Representative structure of the CDK2: carbaryl complex after a 500 ns MD simulation. The inset shows key residues of CDK2 involved in interactions with carbaryl throughout the 500 ns simulation. **(B)** RMSD versus radius of gyration (Rg) plot of CDK2 in the apo (green) and carbaryl-bound (red) forms, demonstrating the structural stability and compactness of CDK2 throughout the 500 ns MD simulation. **(C)** Root mean square fluctuation (RMSF) of CDK2 residues in the apo (green) and carbaryl-bound (wheat) forms. **(D)** Delta RMSF (ΔRMSF) for individual CDK2 residues (nm), illustrating changes in residue fluctuations. **(E)** The zoomed view of the RMSF of the G-rich loop (residues 15-24), C-loop (residues 45-56), hinge loop (residues 80-90), activation loop (residues 152-165), and C-loop substrate recognition (residues 205-217) of CDK2 in the apo (green) and carbaryl-bound (wheat) forms. **(F)** Delta relative solvent-accessible surface area (ΔrSASA) expressed as a percentage for individual CDK2 residues, illustrating changes in solvent accessibility. **(G)** Two-dimensional overlay and change in free energy landscape (ΔFEL) of CDK2 in the apo and carbaryl-bound forms. The global minimum energy state in the apo form is indicated by a filled circle (●) and in the carbaryl-bound form by a filled star (★). The MD simulations and the corresponding analyses were performed in triplicate, and the representative graphs show the mean across three independent simulations for each parameter analysed.

CETSA confirmed direct binding of carbaryl to both kinases in cells, but with opposite thermal signatures: a stabilising shift (increased T_agg_) for CDK1 and a destabilising shift (decreased T_agg_) for CDK2, mirroring the MD predictions **(Figure 6A-D)**. Functionally, *in vitro* kinase activity was significantly reduced for CDK1 but unchanged for CDK2 **(Figure 6E)**.

**Figure 6.**
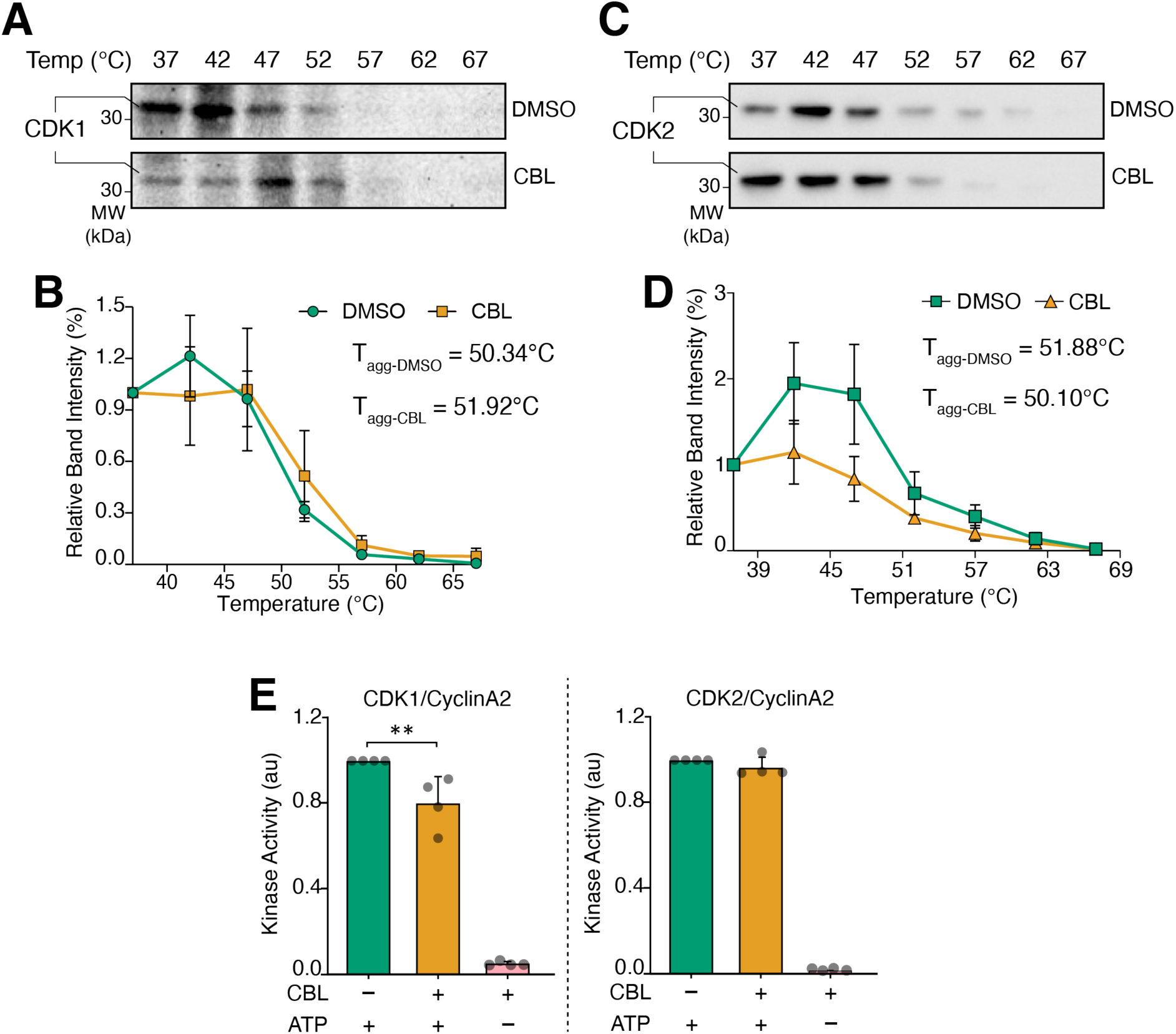
Carbaryl directly targets CDK1 and CDK2 but only modulates CDK1 kinase activity. **(A)** IMCD3 cells were treated with DMSO or carbaryl for 1 h, followed by a 3-minute incubation at temperatures ranging from 37 ℃ to 67 ℃ in 5 ℃ increments. Cells were lysed, and the lysates were immunoblotted for CDK1. **(B)** Quantification of CDK1 protein stability. Signal intensities for CDK1 were measured and plotted. CDK1 expression at 37 ℃ in both conditions was set to 100%. Data represent the mean ±SD of four independent experiments. **(C)** IMCD3 cells were treated with DMSO or carbaryl for 1 h, followed by a 3-minute incubation at temperatures ranging from 37 ℃ to 67 ℃ in 5 ℃ increments. Cells were lysed, and the lysates were immunoblotted for CDK2. **(D)** Quantification of CDK2 protein stability. Signal intensities for CDK2 were measured and plotted. CDK2 expression at 37 ℃ in both conditions was set to 100%. Data represent the mean ±SD of four independent experiments. **(E)** Quantification of *in vitro* CDK1/CyclinA2 and CDK2/CyclinA2 kinase activity in the presence of carbaryl. Kinase activity in the absence of carbaryl was set to 1. Data represent the mean ± SD of four independent experiments. Kinase activity is expressed in arbitrary units (a.u.). Asterisks denote statistical significance as determined by unpaired Student’s t-test: **** p ≤ 0.0001; *** p ≤ 0.001; ** p ≤ 0.01; * p ≤ 0.05.

## Discussion

Carbaryl is a recognised cholinesterase inhibitor that also induces oxidative stress, endocrine disruption, and carcinogenic N-nitroso derivative formation [11,20–22], yet its cellular mechanism remains unclear. Screening across cell lines revealed differential toxicity restricted to IMCD3 and HeLa cells, prompting us to evaluate *Cyp1a1/1a2* expression, since CYPs are critical for carbaryl metabolism [19]. Baseline *Cyp1a1* was lower in IMCD3 than in NIH3T3 and decreased further upon treatment, counter to the classical AhR-mediated induction seen with many xenobiotics, and we cannot yet distinguish cause from consequence of arrest.

The coordinated increase in cell and nuclear size with a preserved nucleus-to-cytoplasm ratio is a hallmark of cellular hypertrophy accompanying G2/M arrest, consistent with metabolically active arrest rather than growth cessation. Increased mitochondrial mass without a rise in ROS reinforces that carbaryl-induced apoptosis is ROS-independent, arguing against classical oxidative-stress-mediated genotoxicity. The hyperchromic shift in DNA absorbance accompanying binding is distinct from the hypochromism classically associated with intercalation, more consistent with helix destabilisation or groove binding [23]; this occurred with parent carbaryl, absent nitrosating agents, indicating a genotoxic route distinct from the previously described nitrosation-dependent mechanism.

The apoptotic phenotype is informative alongside the *Bcl-2* data. *Bcl-2* is canonically anti-apoptotic, yet its upregulation here coincided with an increased late-apoptotic, rather than early-apoptotic, population, suggesting a failed or delayed compensatory survival attempt by cells already committed to death following prolonged checkpoint arrest. This is consistent with reports that sustained mitotic arrest engages slow, *Bcl-2*-family-regulated intrinsic apoptosis via mitotic slippage or catastrophe, and aligns with our finding that carbaryl-induced mitotic arrest in IMCD3 is irreversible within our observation window after carbaryl withdrawal [24–26].

CDK1 and CDK2 share over 60% sequence identity but have distinct roles: CDK2 is dispensable for somatic interphase progression; CDK1 compensates [27–31], whereas CDK1 is essential for viability [32]. Multiple lines of evidence converge on CDK1 as a direct, functionally consequential target: MD simulations predict destabilisation of the G-loop, hinge, and substrate-recognition loop; CETSA shows a stabilising thermal shift consistent with direct pocket engagement; and in vitro kinase activity is significantly reduced. We propose that carbaryl engages CDK1 directly at the ATP-binding cleft to reduce its kinase activity; a mechanism consistent with the literature to produce G2/M arrest. By contrast, carbaryl engages CDK2 through an opposing, rigidifying mode with no change in abundance or activity, an asymmetry that coherently explains the combined S-phase and G2/M arrest.

These findings offer a plausible cellular mechanism consistent with several genotoxic, tissue-selective outcomes reported in human epidemiology, rather than any single clinical phenotype: chronic, sublethal stress driving arrest and apoptosis rather than immediate death is compatible with elevated cancer risk and chromosomal aberration/micronuclei frequency in exposed cohorts, and, in renal epithelium specifically, with the gradual tubular loss characterising CKDu, unlike acute necrosis from classical nephrotoxicants. We emphasise these are candidate correlates from published human data, not in vivo validation: we did not assess renal, oncogenic, or reproductive phenotypes in animals, so the mechanism is hypothesis-generating rather than direct evidence for any one outcome. Two caveats apply: CKDu is a proximal tubular nephropathy while IMCD3 derives from the distal collecting duct, so we regard it as representative rather than segment-matched; and in vitro concentrations (EC50/IC50 23-129 µM, 48 h) exceed typical chronic exposures, so real-world relevance remains to be established.

In summary, carbaryl disrupts cell cycle progression in a lineage-selective manner, most severely in IMCD3 cells, through direct DNA binding, ROS-independent *Bcl-2*-mediated late apoptosis, irreversible mitotic arrest, and CDK1 modulation, establishing it as a genotoxic, cell-cycle-disrupting agent with tissue-selective potency relevant to its reported associations with renal and other tissue pathologies in humans.

## Materials and Methods

### Reagents and tools

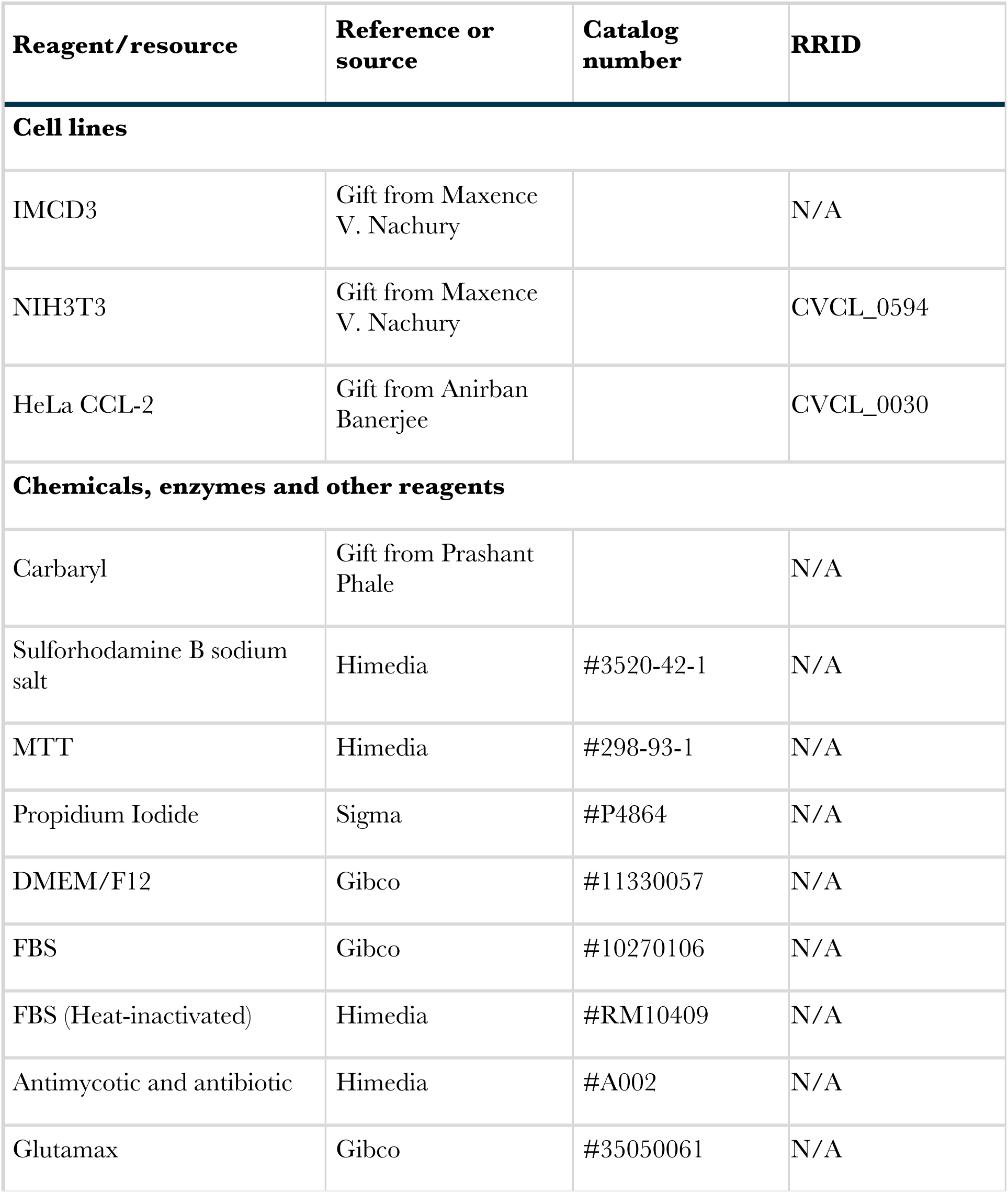
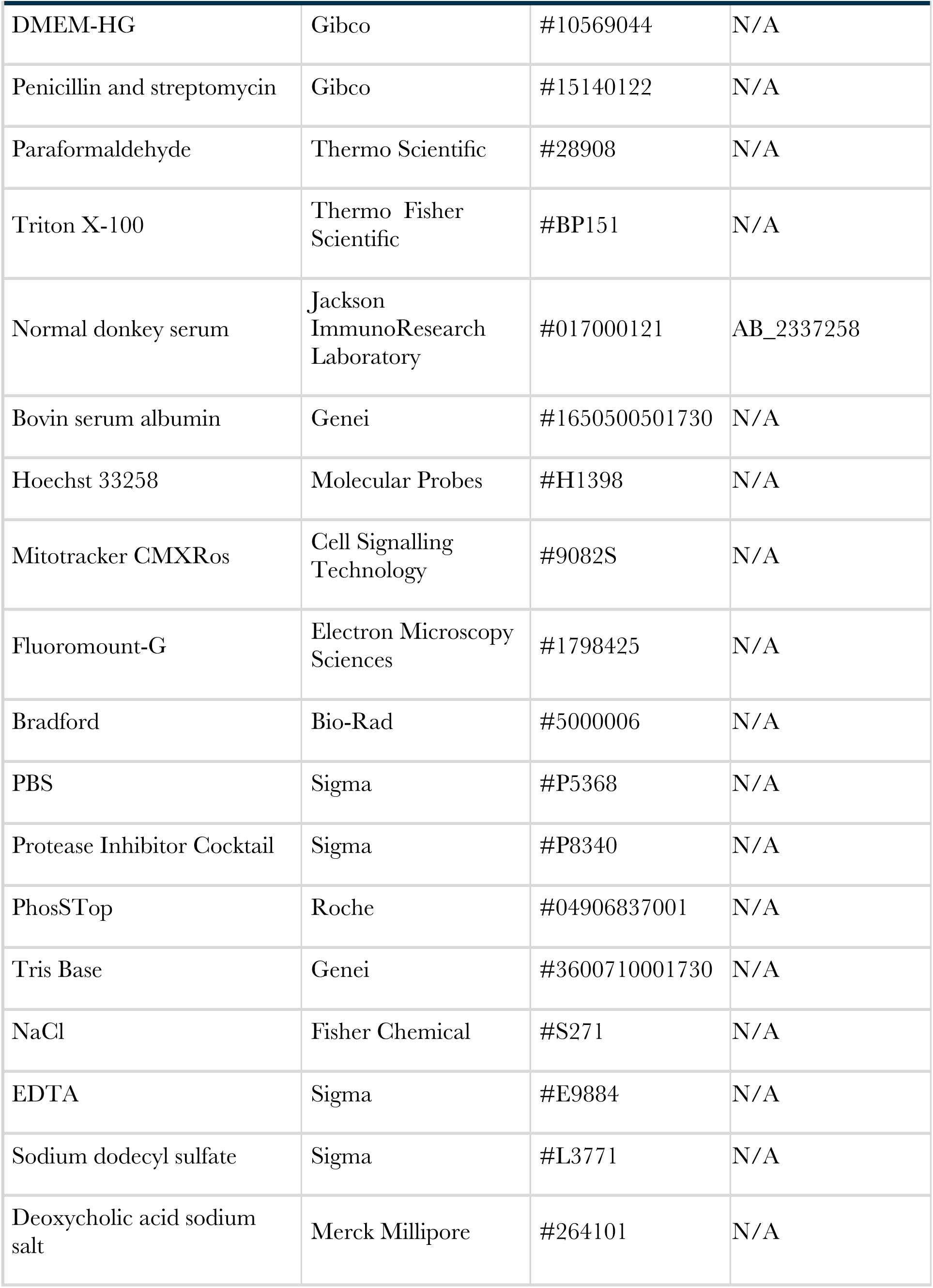
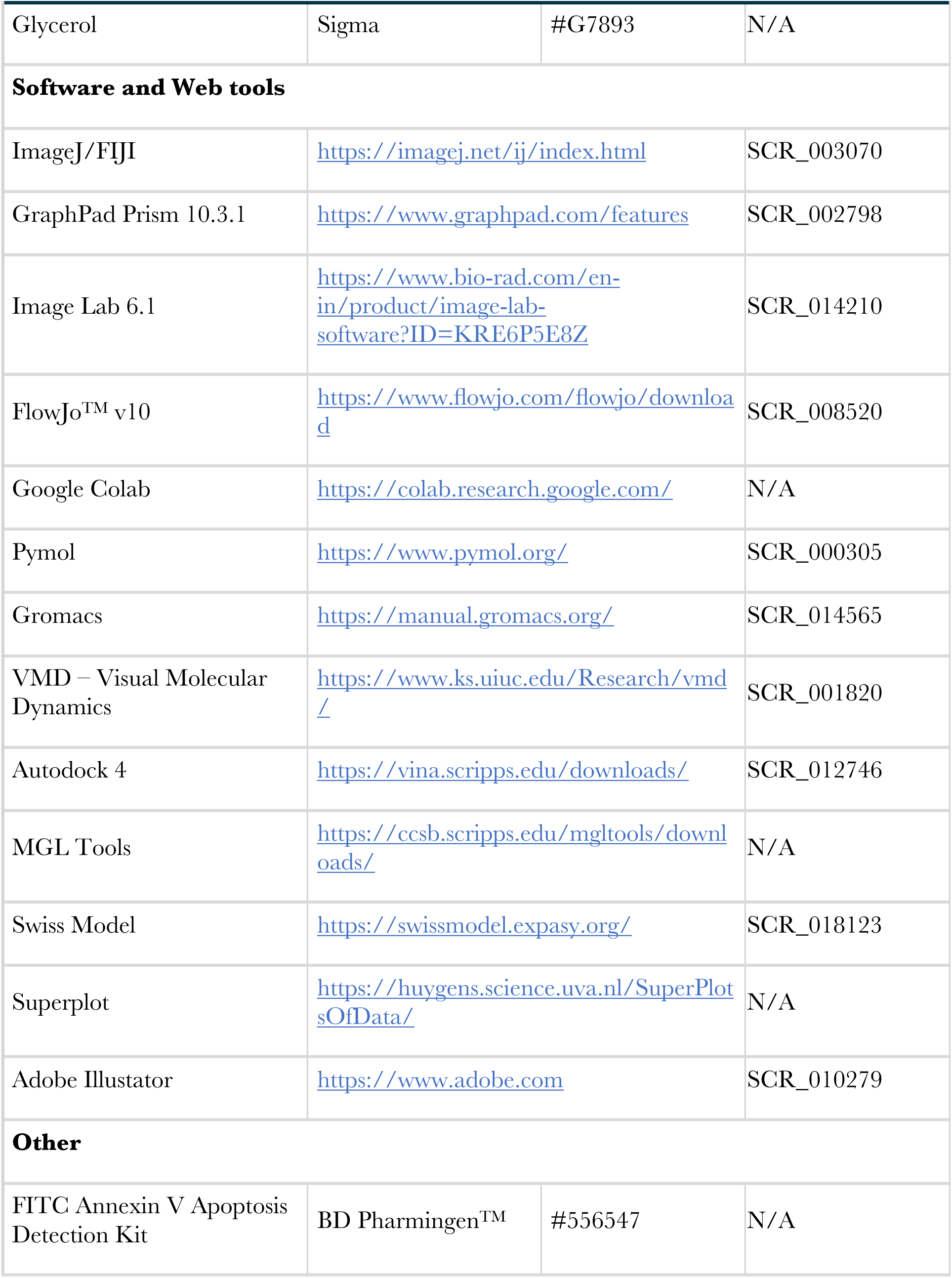
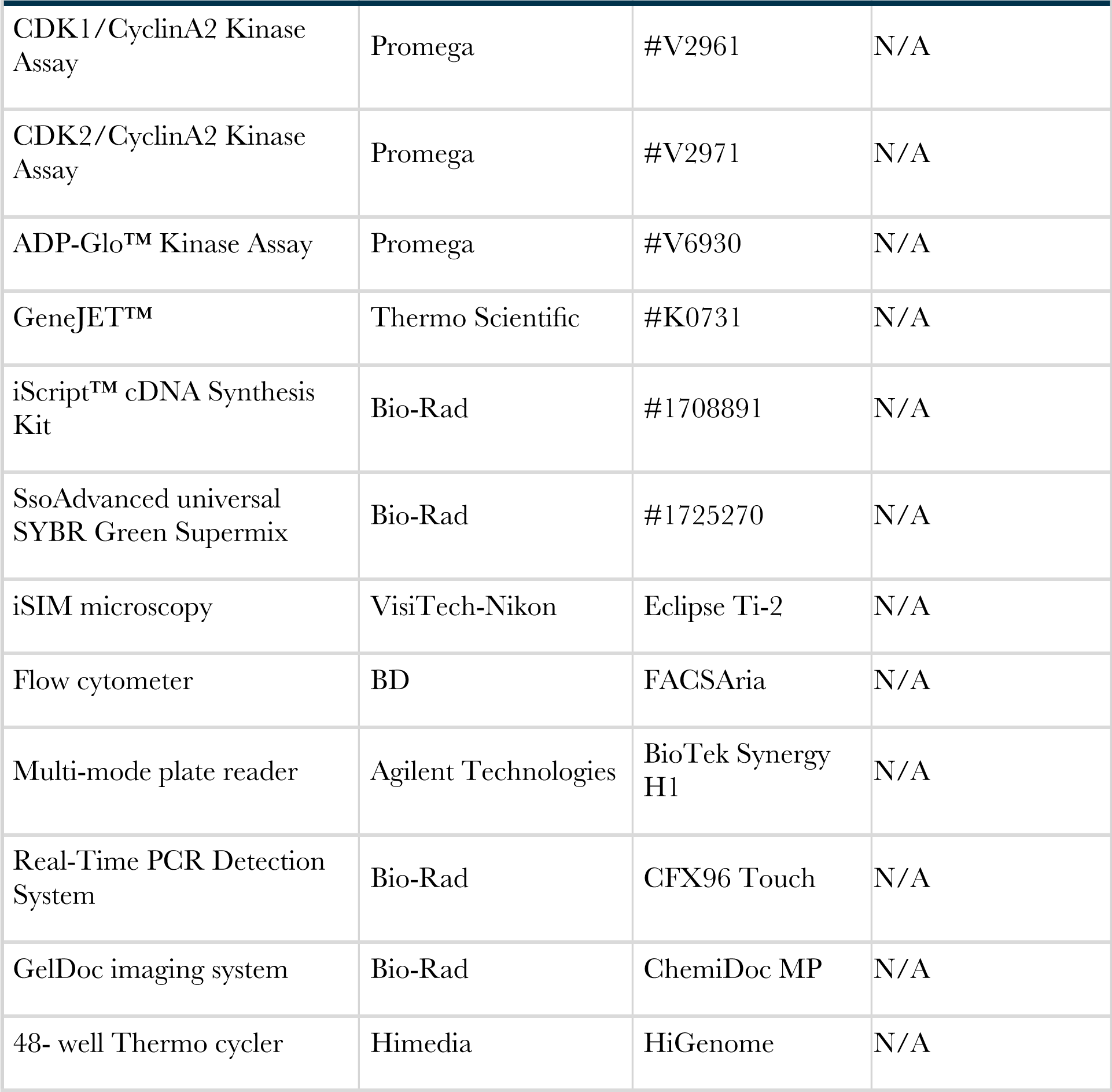

**Reagents and tools:**
| Reagent/resource | Reference or source | Catalog number | RRID |
| --- | --- | --- | --- |
| **Cell lines** |  |  |  |
| IMCD3 | Gift from Maxence V. Nachury |  | N/A |
| NIH3T3 | Gift from Maxence V. Nachury |  | CVCL_0594 |
| HeLa CCL-2 | Gift from Anirban Banerjee |  | CVCL_0030 |
| **Chemicals, enzymes and other reagents** |  |  |  |
| Carbaryl | Gift from Prashant Phale |  | N/A |
| Sulforhodamine B sodium salt | Himedia | #3520-42-1 | N/A |
| MTT | Himedia | #298-93-1 | N/A |
| Propidium Iodide | Sigma | #P4864 | N/A |
| DMEM/F12 | Gibco | #11330057 | N/A |
| FBS | Gibco | #10270106 | N/A |
| FBS (Heat-inactivated) | Himedia | #RM10409 | N/A |
| Antimycotic and antibiotic | Himedia | #A002 | N/A |
| Glutamax | Gibco | #35050061 | N/A |

| **Reagent/resource** | **Reference or source** | **Catalog number** | **RRID** |
| --- | --- | --- | --- |
| DMEM-HG | Gibco | #10569044 | N/A |
| Penicillin and streptomycin | Gibco | #15140122 | N/A |
| Paraformaldehyde | Thermo Scientific | #28908 | N/A |
| Triton X-100 | Thermo Fisher Scientific | #BP151 | N/A |
| Normal donkey serum | Jackson ImmunoResearch Laboratory | #017000121 | AB_2337258 |
| Bovin serum albumin | Genei | #1650500501730 | N/A |
| Hoechst 33258 | Molecular Probes | #H1398 | N/A |
| Mitotracker CMXRos | Cell Signalling Technology | #9082S | N/A |
| Fluoromount-G | Electron Microscopy Sciences | #1798425 | N/A |
| Bradford | Bio-Rad | #5000006 | N/A |
| PBS | Sigma | #P5368 | N/A |
| Protease Inhibitor Cocktail | Sigma | #P8340 | N/A |
| PhosSTop | Roche | #04906837001 | N/A |
| Tris Base | Genei | #3600710001730 | N/A |
| NaCl | Fisher Chemical | #S271 | N/A |
| EDTA | Sigma | #E9884 | N/A |
| Sodium dodecyl sulfate | Sigma | #L3771 | N/A |
| Deoxycholic acid sodium salt | Merck Millipore | #264101 | N/A |
| Glycerol | Sigma | #G7893 | N/A |
| **Software and Web tools** |  |  |  |
| ImageJ/FIJI | https://imagej.net/ij/index.html |  | SCR_003070 |
| GraphPad Prism 10.3.1 | https://www.graphpad.com/features |  | SCR_002798 |
| Image Lab 6.1 | https://www.bio-rad.com/en-in/product/image-lab-software?ID=KRE6P5E8Z |  | SCR_014210 |
| FlowJo™ v10 | https://www.flowjo.com/flowjo/download |  | SCR_008520 |
| Google Colab | https://colab.research.google.com/ |  | N/A |
| Pymol | https://www.pymol.org/ |  | SCR_000305 |
| Gromacs | https://manual.gromacs.org/ |  | SCR_014565 |
| VMD – Visual Molecular Dynamics | https://www.ks.uiuc.edu/Research/vmd/ |  | SCR_001820 |
| Autodock 4 | https://vina.scripps.edu/downloads/ |  | SCR_012746 |
| MGL Tools | https://ccsb.scripps.edu/mgltools/downloads/ |  | N/A |
| Swiss Model | https://swissmodel.expasy.org/ |  | SCR_018123 |
| Superplot | https://huygens.science.uva.nl/SuperPlotsOfData/ |  | N/A |
| Adobe Illustrator | https://www.adobe.com |  | SCR_010279 |
| **Other** |  |  |  |
| FITC Annexin V Apoptosis Detection Kit | BD Pharmingen™ | #556547 | N/A |
| CDK1/CyclinA2 Kinase Assay | Promega | #V2961 | N/A |
| CDK2/CyclinA2 Kinase Assay | Promega | #V2971 | N/A |
| ADP-Glo™ Kinase Assay | Promega | #V6930 | N/A |
| GeneJET™ | Thermo Scientific | #K0731 | N/A |
| iScript™ cDNA Synthesis Kit | Bio-Rad | #1708891 | N/A |
| SsoAdvanced universal SYBR Green Supermix | Bio-Rad | #1725270 | N/A |
| iSIM microscopy | VisiTech-Nikon | Eclipse Ti-2 | N/A |
| Flow cytometer | BD | FACSAria | N/A |
| Multi-mode plate reader | Agilent Technologies | BioTek Synergy H1 | N/A |
| Real-Time PCR Detection System | Bio-Rad | CFX96 Touch | N/A |
| GelDoc imaging system | Bio-Rad | ChemiDoc MP | N/A |
| 48- well Thermo cycler | Himedia | HiGenome | N/A |

### Antibodies used in the study

**Antibodies used in the study.**
| Antibody | Species | Dilution | Company | Cat. No. | RRID | Application |
| --- | --- | --- | --- | --- | --- | --- |
| Monoclonal Primary |  |  |  |  |  |  |
| α-tubulin | Mouse | 1:5000<br>and<br>1:1000 | Sigma | T6199 | AB_477583 | IF and WB |
| Acetylated tubulin | Mouse | 1:10000 | Sigma | T7451 | AB_609894 | IF |
| γ-tubulin | Mouse | 1:5000<br>and<br>1:1000 | Sigma | T6557 | AB_477584 | IF and WB |
| β-actin | Mouse | 1:1000 | Santa Cruz<br>Biotechnology | Sc-47778 | AB_626632 | WB |
| γ-H2A.X | Rabbit | 1:500<br>and<br>1:1000 | Cell Signaling<br>Technology | 9718S | AB_2118009 | IF and WB |
| CDK1<br>(cdc2) | Mouse | 1:1000 | Cell Signaling<br>Technology | 9116T | N/A | WB |
| CDK2 | Rabbit | 1:1000 | Cell Signaling<br>Technology | 2546S | AB_2276129 | WB |
| Polyclonal Primary |  |  |  |  |  |  |
| γ-tubulin | Rabbit | 1:500 | Abcam | AB1131<br>7 | AB_297920 | IF |
| Phospho-Histone<br>H3 | Rabbit | 1:1000 | Cell Signaling<br>Technology | 9701L | AB_331535 | IF |
| Fluorophore-tagged secondary |  |  |  |  |  |  |
| Cy5 | Goat | 1:500 | Jackson<br>ImmunoResearch<br>Laboratory | 115-175-205 | N/A | IF |
| Cy5 | Goat |  | Jackson<br>ImmunoResearch<br>Laboratory | 115-175-206 | N/A | IF |
| Cy5 | Donkey |  | Jackson<br>ImmunoResearch<br>Laboratory | 711-175-152 | AB_2340607 | IF |
| Cy3 | Goat |  | Jackson<br>ImmunoResearch<br>Laboratory | 115-165-207 | N/A | IF |
| Cy3 | Goat |  | Jackson ImmunoResearch Laboratory | 115-165-206 | N/A | IF |
| Cy3 | Donkey |  | Jackson ImmunoResearch Laboratory | 711-165-152 | N/A | IF |
| Alexa 488 | Goat |  | Invitrogen | A11001 | AB_2534069 | IF |
| Alexa 488 | Goat |  | Invitrogen | A11008 | AB_143165 | IF |
| HRP-tagged secondary |  |  |  |  |  |  |
| Anti-mouse | Goat | 1:10000 | Cell Signaling Technology | 7076S | AB_330924 | WB |
| Anti-rabbit | Goat |  | Cell Signaling Technology | 7074S | AB_2099233 | WB |

### Oligos used in this study

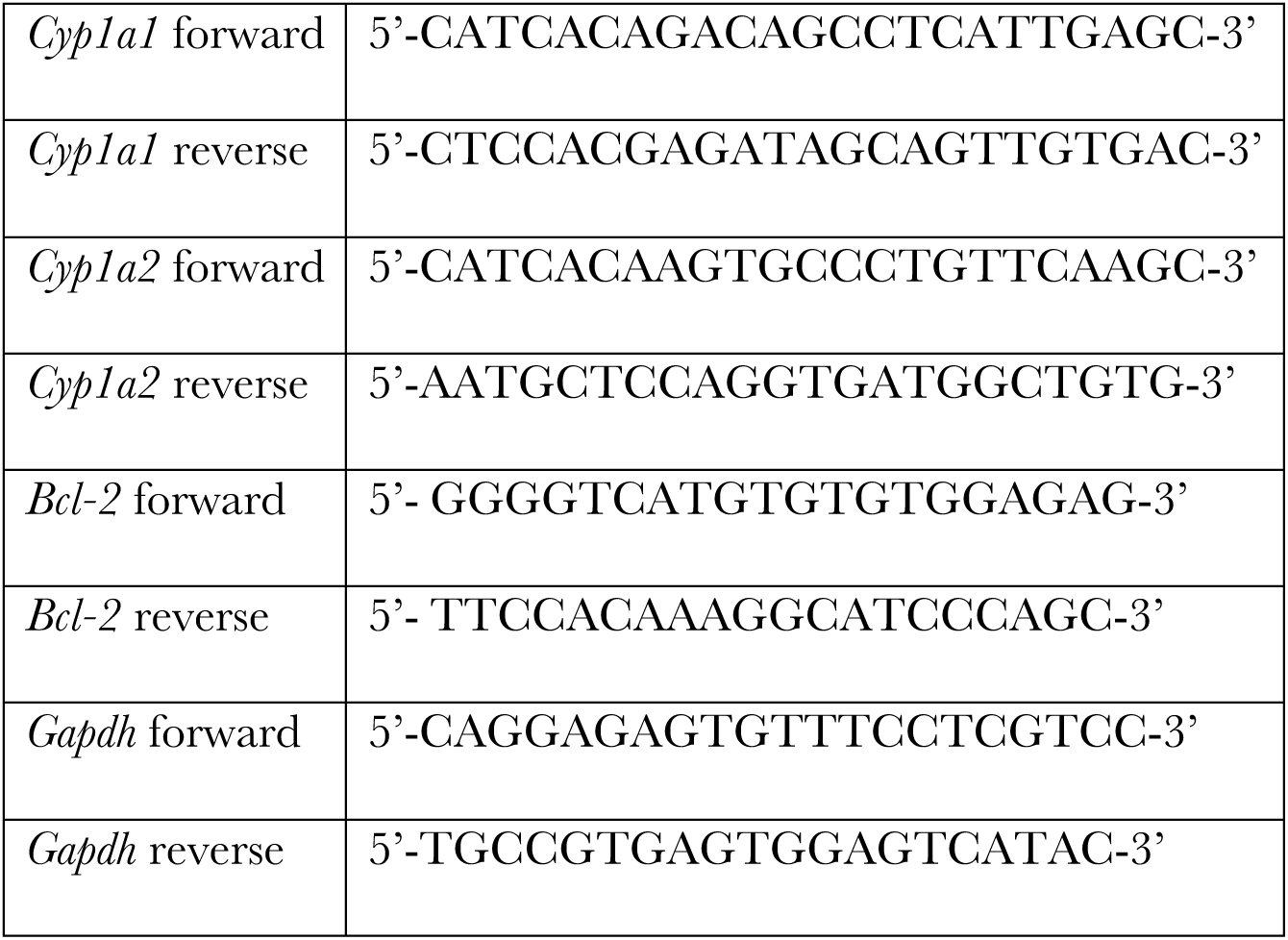

### Cell Culture

The mouse IMCD3-FlpIn cell line (gift from Maxence V. Nachury, UCSF, CA) cells were cultured in DMEM/F12 (Cat. No. 11330057; Gibco) supplemented with 10% FBS (Cat. No. 10270106; Gibco), 1x antimycotic and antibiotic (Cat. No. A002; Himedia), and 1 % Glutamax (Cat. No. 35050061; Gibco). HeLa (kind gift from Anirban Banerjee, IIT Bombay) were cultured in high-glucose DMEM (Cat. No. 10566016; Gibco) supplemented with 10 % FBS and 1% antibiotic-antimycotic solution (Cat. No. A002A; HiMedia). NIH3T3 cells (gift from Maxence V. Nachury, UCSF, CA) were maintained in high-glucose DMEM and supplemented with 10% heat-inactivated FBS (Cat. No. F4135; HiMedia), 1% Pen-strep (Cat. No. 15140122; Gibco). All the cells were incubated at 37 °C with 5% CO_2_.

### Antibodies and drug

The following primary antibodies were used for immunofluorescence: Monoclonal: anti-acetylated tubulin (mouse; clone 6-11B-1; Cat. No. T7451; Sigma-Aldrich; 1:10,000), anti-alpha tubulin (mouse; clone DM1A; Cat. No. T6199; Sigma-Aldrich; 1:5,000), anti-γ-tubulin (mouse; Cat. No. T6557; Sigma-Aldrich; 1:5,000), Phospho-Histone H2A.X (Ser139) (20E3) (rabbit; Cat. No. 9718; Cell Signaling Technology; 1:1000) (kind gift from Shamik Sen, IIT Bombay). Polyclonal: anti-γ-tubulin (Rabbit; AB11317; Abcam; 1:500), phospho-Histone H3 (rabbit; Cat. No. 9701; Cell Signaling Technology; 1:1000). The following dyes were used: MitoTracker Red CMXRos (9082S; Cell Signaling Technology; 200 ng) (kind gift from Roop Malik, IIT Bombay) and Hoechst 33258 (H1398; Molecular Probes). The following fluorophore-tagged secondary antibodies from Jackson ImmunoResearch Laboratory were used: Cy5 anti-mouse IgG1 (goat; Cat. No. 115-175-205), Cy3 anti-mouse IgG2b (goat; Cat. No. 115-165-207), Cy5 anti-rabbit IgG (donkey; Cat. No. 711-175-152). The following fluorophore-tagged secondary antibodies from Invitrogen were used: Alexa 488 anti-mouse (goat; Cat. No. A11001) and Alexa 488 anti-rabbit (goat; Cat. No. A11008). All secondary antibodies were used in 1:500 dilutions.

Carbaryl was a kind gift from Prasant Phale, IIT Bombay. Doxorubicin was a kind gift from Ashutosh Kumar, IIT Bombay.

### Calculation of half-maximal effective and inhibitory concentrations

Cells were seeded in 96-well plates (5000–10,000 cells/well), allowed to attach for 12 h, and treated with carbaryl for 48 h. EC50 was determined by MTT assay: MTT was added (1:9 with media) and incubated for 4 h; formazan crystals were dissolved in 70 µL DMSO (dark, shaking, 35–45 min), and absorbance was read at 570 nm. IC50 was determined by SRB assay: cells were fixed with 30 µL 50% TCA (4 °C, 1 h), washed with water and dried, stained with 100 µL SRB (1 h, dark), rinsed with 1% acetic acid, dried, and dissolved in Tris-base (pH 10.5, 30 min); absorbance was read at 520 nm. Both assays were performed in at least three independent experiments.

### Flow cytometry

For cell cycle analysis, IMCD3-FlpIn and HeLa cells were treated with vehicle (0.05% DMSO) or carbaryl (12 µM and 64 µM, respectively) for 48 h, trypsinised, washed, and fixed in ice-cold 70% ethanol in 1x PBS with 0.1% FBS (−20 °C, overnight). Cells were then washed twice with 1x PBS and stained with 50 µg/mL propidium iodide (Cat. No. P4864) and 1 µg/mL RNase A for 15 min at room temperature before acquisition.

For the apoptosis assay, IMCD3 cells were treated with vehicle or 12 µM carbaryl for 48 h, trypsinised, and processed using the BD Pharmingen™ FITC Annexin V Apoptosis Detection Kit (Cat. No. 556547) per the manufacturer’s protocol. Both assays were acquired on a BD FACSAria flow cytometer (BD Biosciences) and analysed using FlowJo software.

### DCF-DA Assay

For ROS measurement, IMCD3 cells were treated with vehicle (0.05% DMSO), 12 µM carbaryl, or 18 nM doxorubicin (positive control) for 48 h, trypsinised, and collected in PBS. Cells were incubated with 10 µM DCFDA (kind gift from Rahul Purwar, IIT Bombay) for 20 min at 37 °C, centrifuged (1000 x g, 4 min, room temperature), and resuspended in PBS. Fluorescence was measured (excitation 485 nm, emission 520 nm) using a multimode plate reader (Agilent), and carbaryl-treated values were normalised to DMSO for relative ROS levels.

### Immunofluorescence and Imaging

Cells (80,000) were seeded on ethanol-sterilized coverslips (12 mm, No. 1.5; Cat. No. 12-545-81; Thermo Fisher Scientific) and allowed to adhere for 12 h (37 °C, 5% CO₂). Cells were treated with vehicle (0.05% DMSO) or 12 µM carbaryl for 48 h in complete medium, followed by either no further treatment (condition 1), 16 h serum starvation (condition 2), or 16 h in complete medium (condition 3).

Cells were fixed with 4% paraformaldehyde (Cat. No. 28908; Thermo Scientific) in PBS (15 min, 37 °C), washed twice with PBS, extracted in ice-cold methanol (−20 °C, 5 min), and washed three times with PBS. Cells were blocked/permeabilized in IF buffer (PBS with 0.1% Triton X-100 [Cat. No. BP151; Thermo Fisher Scientific], 5% normal donkey serum [Cat. No. 017-000-121; Jackson ImmunoResearch], 3% BSA [Cat. No. 1650500501730; Genei]) for 30 min, then incubated overnight at 4 °C with primary antibodies in IF buffer. After three IF-buffer washes, cells were incubated with secondary antibodies (Jackson ImmunoResearch) in IF buffer for 30 min at room temperature, then washed three times. DNA was stained with Hoechst 33258 (H1398; Molecular Probes) for 4 min and washed twice with PBS. Coverslips were mounted with Fluoromount-G (17984-25; Electron Microscopy Sciences) and cured before imaging.

Imaging was performed on an instant structured illumination microscope (iSIM; VT-iSIM, Visitech International, on a Nikon Ti2, Nikon Instruments) using a 60x Plan-Apochromat 1.4 NA oil immersion objective, with Z-stacks acquired at 0.5 µm intervals.

### Microtubule Cold Shock Recovery Assay

For the microtubule recovery assay, IMCD3-FlpIn, NIH3T3, and HeLa cells were treated with either the vehicle control (0.05% DMSO) or carbaryl (12 µM for IMCD3-FlpIn and 64 µM for NIH3T3 and HeLa cells) for 48 h. Post-treatment, the cells were subjected to cold shock using cold media and followed by incubation at 4 °C for 1 h to depolymerise the tubulin network. After 1 h, the tubulin network was allowed to regrow at 37 °C for 0, 1, 2, 5, and 10 mins. Cells were washed with PBS and fixed with methanol (pre-chilled at −20 °C) for 10 mins at −20 °C. Immunofluorescence staining for alpha-tubulin and imaging were done as described earlier.

### Western Blotting and Densitometry

IMCD3 lysates were prepared in RIPA buffer (50 mM Tris-HCl pH 7.5, 150 mM NaCl, 1 mM EDTA, 1% Triton X-100, 0.1% SDS, 1% sodium deoxycholate) with freshly added phosphatase/protease inhibitors, passed through a Hamilton syringe (51 mm), and incubated on ice for 20 min. Debris was removed by centrifugation (21,000 x g, 20 min, 4 °C) and the supernatant retained.

Protein concentration was determined by Bradford assay (BSA standard in RIPA buffer). Samples were denatured (95 °C, 5 min, SDS loading buffer) and 30 µg protein/lane resolved by SDS-PAGE. Immunoblotting used the following primary antibodies (1:1000): anti-phospho-Histone H2A.X (Ser139) (20E3; rabbit; Cat. No. 9718; Cell Signaling Technology; gift from Shamik Sen, IIT Bombay), anti-alpha tubulin (mouse; clone DM1A; Cat. No. T6199; Sigma-Aldrich; 1:1,000), anti-γ-tubulin (mouse; Cat. No. T6557; Sigma-Aldrich; 1:1,000), and anti-β-actin (clone C4; mouse; Cat. No. sc-47778; Santa Cruz Biotechnology).

HRP-linked secondary antibodies (1:10,000) were anti-mouse IgG (Cat. No. 7076S) and anti-rabbit IgG (Cat. No. 7074S; both Cell Signaling Technology). Blots were developed on a ChemiDoc MP imaging system (Bio-Rad).

Blots were quantified in ImageLab 6.1 (Bio-Rad); band intensity for proteins of interest was normalised to β-actin (loading control) to compare protein levels across conditions.

### Cellular Thermal Shift Assay

IMCD3-FlpIn cells (80–90% confluent, 10 cm dishes) were treated with carbaryl for 1 h, trypsinized, washed, and resuspended in PBS. Aliquots (1.5 × 10⁶ cells) were heated across a 37–67 °C gradient (5 °C steps, 3 min, 48-well thermocycler), then cooled to 25 °C for 3 min. Cells were lysed by three freeze-thaw cycles in liquid nitrogen, and lysates were clarified by centrifugation (10,000 × g, 20 min, 4 °C). Protein concentration was measured by Bradford assay (BSA standards), normalising to the 37 °C and 42 °C samples. Samples were boiled (95 °C, 5 min, loading buffer), and 30 µg protein/lane was resolved by SDS-PAGE and immunoblotted with anti-CDK1 (rabbit; Cat. No. 9116T; Cell Signaling Technology; 1:1000) and anti-CDK2 (rabbit; Cat. No. 2546S; Cell Signaling Technology; 1:1000; gift from Subbareddy Maddika, CDFD, Hyderabad). Blots were visualized on a ChemiDoc MP system (Bio-Rad).

### UV-vis spectroscopy

100 ng of plasmid DNA was incubated in the absence and presence of Carbaryl (12 µM and 64 µM) in 10 mM PBS for 15 min at room temperature. The absorbance spectra were measured from 230-320 nm with a step size of 5 nm using a multimode plate reader (Agilent). The peak at 260 nm was compared to observe the change in DNA spectra in the presence of carbaryl.

### Fluorescence spectroscopy

100 ng of plasmid DNA was incubated in the absence and presence of Carbaryl (12 µM and 64 µM) in 10 mM PBS for 15 min at room temperature. EtBr was kept as a positive control. The fluorescence spectra were measured from 300-700 nm with a step size of 2 nm using a multimode plate reader (Agilent). The peak at 330 nm was compared to observe the change in carbaryl spectra in the presence of DNA.

### qRT-PCR

Total RNA was extracted from IMCD3 and NIH3T3 cells after 48 h of carbaryl treatment (concentrations as in the microtubule cold-shock recovery assay) using the RNASure Mini Kit (Genetix, NP-84105). 1 µg RNA was used for cDNA synthesis (iScript cDNA Synthesis Kit; Cat. No. 1708891; Bio-Rad), per manufacturer’s protocols. qRT-PCR was performed with SsoAdvanced Universal SYBR Green Supermix (Cat. No. 1725270; Bio-Rad) in 10 µL reactions (30 ng cDNA template) on a CFX96 Touch Real-Time PCR Detection System (Bio-Rad), with *Gapdh* as the reference gene. Relative expression was calculated by the 2-ΔΔCt method from technical triplicates. Primers are listed in Table S2.

### Molecular Dynamics Simulation

All-atom MD simulations were performed for apo and carbaryl-bound CDK1 (PDB ID 6GU2, 2 Å), CDK2 (PDB ID 1H1P, 2.1 Å), and CDK4 (PDB ID: 3G33, Resolution: 3 Å), using the highest-energy docking pose, in GROMACS (v2024.4). Protein topology was generated with the CHARMM27 force field [40], and ligand topology with ACPYPE [41]. Systems were solvated in a triclinic box and neutralized with Na⁺/Cl⁻ to 0.15 M. Energy minimization used the steepest descent algorithm and Verlet cut-off scheme (max force tolerance 10 kJ mol⁻¹ nm⁻¹). Equilibration proceeded through sequential NVT and NPT phases (100 ps/50,000 steps each), maintaining 300 K (V-rescale thermostat, 0.1 ps coupling) and 1 bar (Parrinello-Rahman barostat) [42,43], with coordinates saved every 2 ps. Production simulations ran for 500 ns. Trajectories were analysed using GROMACS and VMD; visualizations were rendered in PyMOL [44]. RMSD, Rg, and free energy landscape plots were generated via a custom Python script in Google Colab; other graphs used GraphPad Prism (v10.0). Simulations were run on the Spacetime HPC cluster, IIT Bombay.

### Image analysis

Image files were imported into ImageJ/Fiji (National Institutes of Health, Bethesda, MD, USA) for analysis. For quantification of fluorescence signal intensity in fixed cells, maximum intensity projections (MIPs) were used. For pH3-positive cell count, the same dynamic range was set for treated and untreated cells. Cells showing signal intensity were considered positive. Data were plotted as SuperPlots using the SuperPlotsOfData web application [33]. Each SuperPlot displays the full distribution of individual data points, with the mean value of each independent experiment represented by a larger symbol. All representative micrographs within each panel are displayed using the same dynamic range.

### Statistical analysis

All curve fits for IC_50_ were performed using GraphPad Prism (version 10.3.1). The non-linear regression curve-fit model with log(inhibitor) vs. response - Variable slope (four parameters)- least-squares fit method was chosen. For EC_50_, the non-linear regression curve-fit model with log(agonist) vs. response - Variable slope (four parameters)- least-squares fit method was chosen. For T_agg,_ the non-linear regression curve-fit model with a Boltzmann sigmoidal form was chosen. We used an unpaired Student’s t-test between two experimental groups. For more than two conditions, we performed one-way ANOVA followed by Tukey’s post hoc test to determine significance using GraphPad Prism. In bar graphs and superplots, data are represented as the mean ± SEM. Asterisks indicate statistical significance values calculated by unpaired Student’s t-test and one-way ANOVA followed by Tukey’s post hoc multiple comparison test. p-value ≤ 0.0001; ****, p-value ≤ 0.001; ***, p-value ≤ 0.01; **, p-value ≤ 0.05; *.

## Author contributions

R.P. and S.R.S. designed the experiments; R.P. and S.T. performed all experiments. R.P., S.T., and S.R.S. wrote the manuscript. R.P. and S.R.S. conceptualised the study. S.R.S. supervised the research and secured funding.

## Acknowledgement

We thank Prasant Phale for providing carbaryl; Ashutosh Kumar for doxorubicin; Anirban Banerjee for kindly providing the HeLa cell lines; and Shamik Sen and Subba Reddy Maddika for providing antibodies. We thank the members of the Cilia Biology Lab for valuable scientific discussions and critical reading of the manuscript. Imaging was performed on the iSIM microscope at the Research Infrastructure Facility Centre (RIFC), IIT Bombay (IE/19-RIFC0000-CF-04SRFM). Flow cytometry was performed at the FACS Central Facility, IIT Bombay, supported by DST FIST (SR/FST/LSI-055/2000). Computational resources were provided by the Spacetime High-Performance Computing cluster at IIT Bombay. RP acknowledges a Senior Research Fellowship from the Department of Biotechnology (DBT), Government of India (DBT/2021-22/IIT-B/1661). ST acknowledges a fellowship from IIT Bombay. This work was supported by the following grants to SRS: a seed grant from IIT Bombay (RD/0522-IRCCSHO-026); a Core Research Grant (CRG/2023/002870) from the Science and Engineering Research Board (SERB), Department of Science and Technology, India; and the Department of Biotechnology (Grant No. BT/PR56541/BMS/85/481/2024). The authors used Grammarly for grammar and spell-checking, and Claude (Anthropic) for assistance with manuscript restructuring. All content was reviewed, verified, and approved by the authors, who take full responsibility for the integrity of the work.

## Supplementary Figures

**Figure S1.**
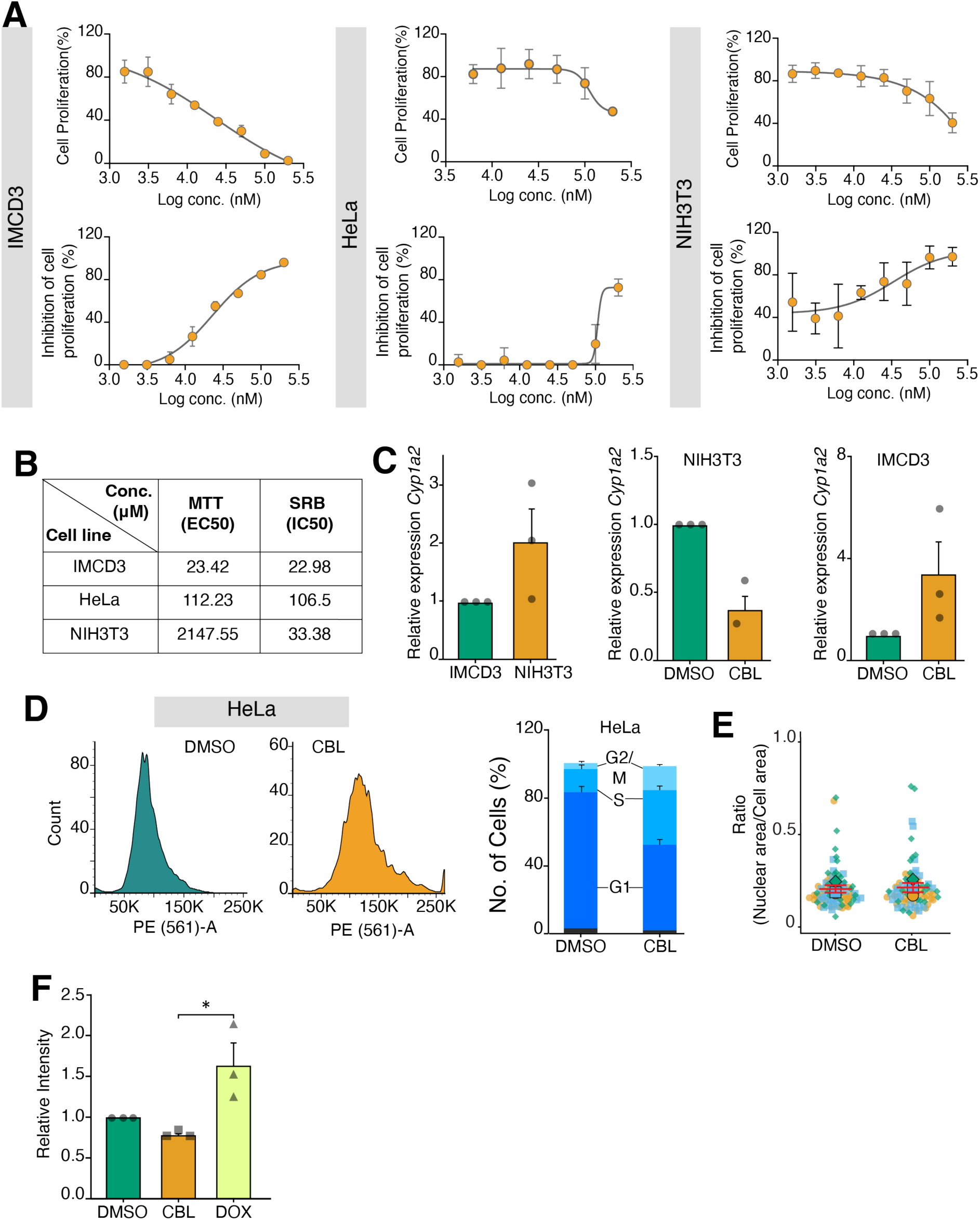
Carbaryl exhibits differential potency. **(A)** Dose-response curves of cell proliferation (top row) and corresponding inhibition of cell proliferation (bottom row) following carbaryl treatment in IMCD3 (left), HeLa (middle), and NIH3T3 (right) cells. **(B)** Table summarizing EC50 (MTT assay) and IC50 (SRB assay) values (µM) for carbaryl across the three cell lines. **(C)** Relative expression of *Cyp1a2* by qPCR. Left: baseline *Cyp1a2* expression is lower in IMCD3 cells compared to NIH3T3 cells. Middle: in NIH3T3 cells, carbaryl (CBL) treatment significantly downregulates *Cyp1a2* expression relative to DMSO control. Right: in IMCD3 cells, carbaryl treatment trends toward increased *Cyp1a2* expression relative to DMSO. Data represent mean ± SEM from three independent experiments (dots represent individual replicates). **(D)** Representative flow cytometry histograms of DNA content in HeLa cells treated with DMSO or carbaryl. Quantification of cell cycle phase distribution in DMSO- versus carbaryl-treated HeLa cells. **(E)** Quantification of the cell area-to-nucleus area ratio in DMSO- versus carbaryl-treated IMCD3 cells. In the superplots, three independent experimental data points are represented by three different shapes, with colours indicated. Each square, circle, or triangle represents a data point, and the larger version of its shape represents the mean. Data represent all data points from three independent experiments, mean ±SEM. DMSO n = 89, carbaryl n = 84. **(F)** Relative fluorescence intensity from a ROS detection assay in DMSO-, carbaryl-, and doxorubicin (positive control)-treated IMCD3 cells. Data represent all data points from three independent experiments, mean ±SEM. Asterisks denote statistical significance as determined by unpaired Student’s t-test or one-way ANOVA followed by Tukey’s post hoc multiple comparison test, as appropriate: **** p ≤ 0.0001; *** p ≤ 0.001; ** p ≤ 0.01; * p ≤ 0.05.

**Figure S2.**
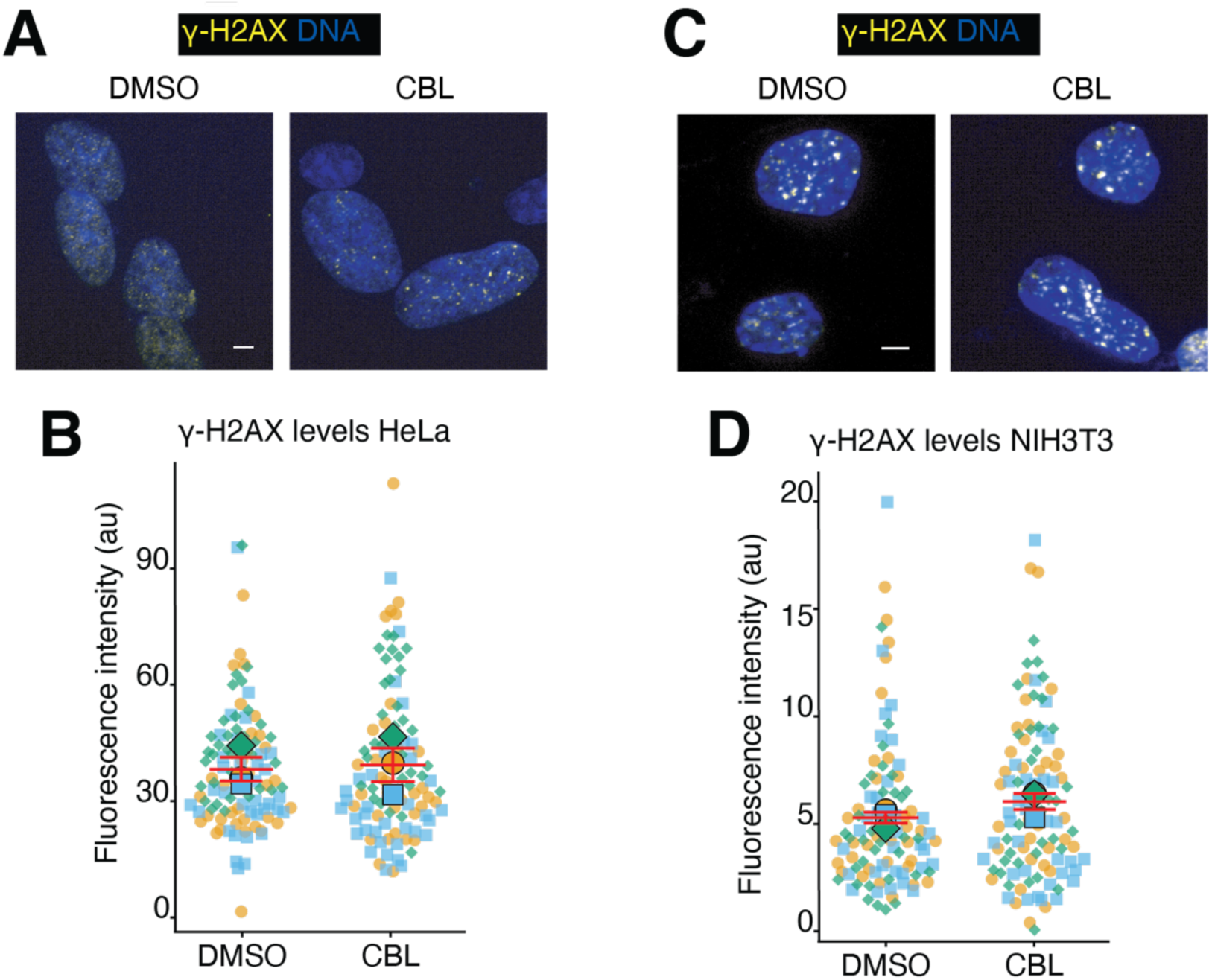
Carbaryl does not induce DNA damage in HeLa or NIH3T3 cells. **(A)** Representative immunofluorescence images of HeLa cells stained for γ-H2AX (yellow) and DNA (blue) following DMSO or carbaryl treatment, showing no visible increase in γ-H2AX foci upon carbaryl exposure. Scale bar: 5 µm. **(B)** Quantification of γ-H2AX fluorescence intensity per nucleus in DMSO- versus carbaryl-treated HeLa cells. In the superplots, three independent experimental data points are represented by three different shapes, with colours indicated. Each square, circle, or triangle represents a data point, and the larger version of its shape represents the mean. Data represent all data points from three independent experiments, mean ±SEM. DMSO n = 105, carbaryl n = 105. **(C)** Representative immunofluorescence images of NIH3T3 cells stained for γ-H2AX (yellow) and DNA (blue) following DMSO or carbaryl treatment. Scale bar: 5 µm. **(D)** Quantification of γ-H2AX fluorescence intensity per nucleus in DMSO- versus carbaryl-treated NIH3T3 cells, showing no significant difference between conditions. Superplot convention is as in **B**. Data represent all data points from three independent experiments, mean ±SEM. DMSO n = 103, carbaryl n = 105. Fluorescence intensities are expressed in arbitrary units (a.u.). Asterisks denote statistical significance as determined by unpaired Student’s t-test: **** p ≤ 0.0001; *** p ≤ 0.001; ** p ≤ 0.01; * p ≤ 0.05.

**Figure S3.**
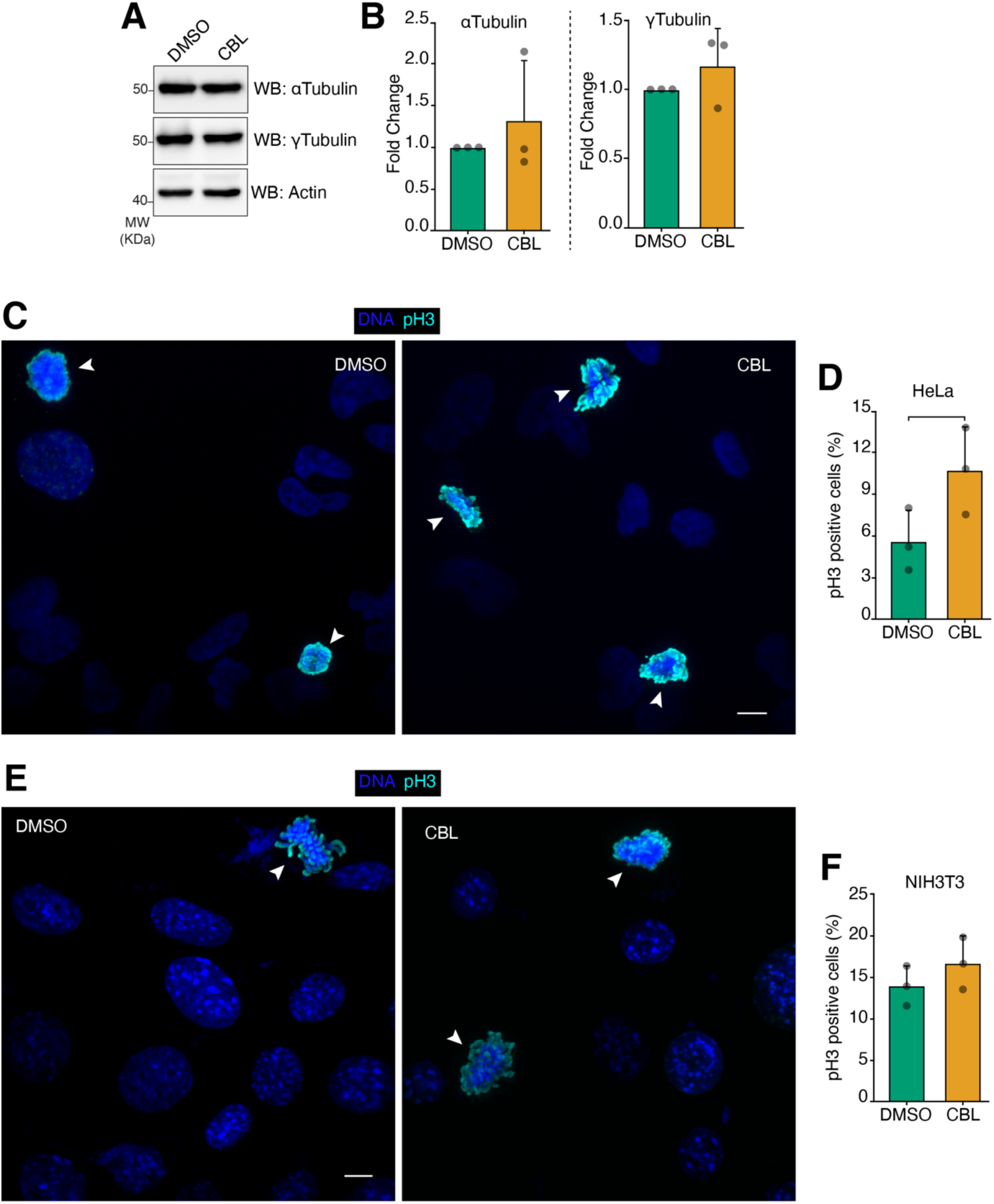
Carbaryl does not alter the mitotic index in HeLa or NIH3T3 cells. **(A)** IMCD3 cells were lysed, and the lysates were probed for alpha-tubulin, gamma-tubulin, and actin (loading control) via immunoblotting. **(B)** Alpha-tubulin and gamma-tubulin signal intensities were measured and normalized against the loading control. Alpha-tubulin and gamma-tubulin expression in the DMSO condition was set to 1. The data represent the mean ±SEM of three independent experiments. **(C)** Representative immunofluorescence images of HeLa cells stained for phospho-histone H3 (pH3, cyan) and DNA (blue) following DMSO or carbaryl treatment. White arrowheads indicate pH3-positive cells. Scale bar: 10 µm. **(D)** Quantification of the percentage of pH3-positive (mitotic) cells in DMSO- versus carbaryl-treated HeLa cells. Data represent the mean ± SEM from three independent experiments. **(E)** Representative immunofluorescence images of NIH3T3 cells stained for phospho-histone H3 (pH3, cyan) and DNA (blue) following DMSO or carbaryl treatment. White arrowheads indicate pH3-positive cells. Scale bar: 10 µm. **(F)** Quantification of the percentage of pH3-positive (mitotic) cells in DMSO- versus carbaryl-treated NIH3T3 cells. Data represent the mean ± SEM from three independent experiments. Asterisks denote statistical significance as determined by unpaired Student’s t-test: **** p ≤ 0.0001; *** p ≤ 0.001; ** p ≤ 0.01; * p ≤ 0.05.

**Figure S4.**
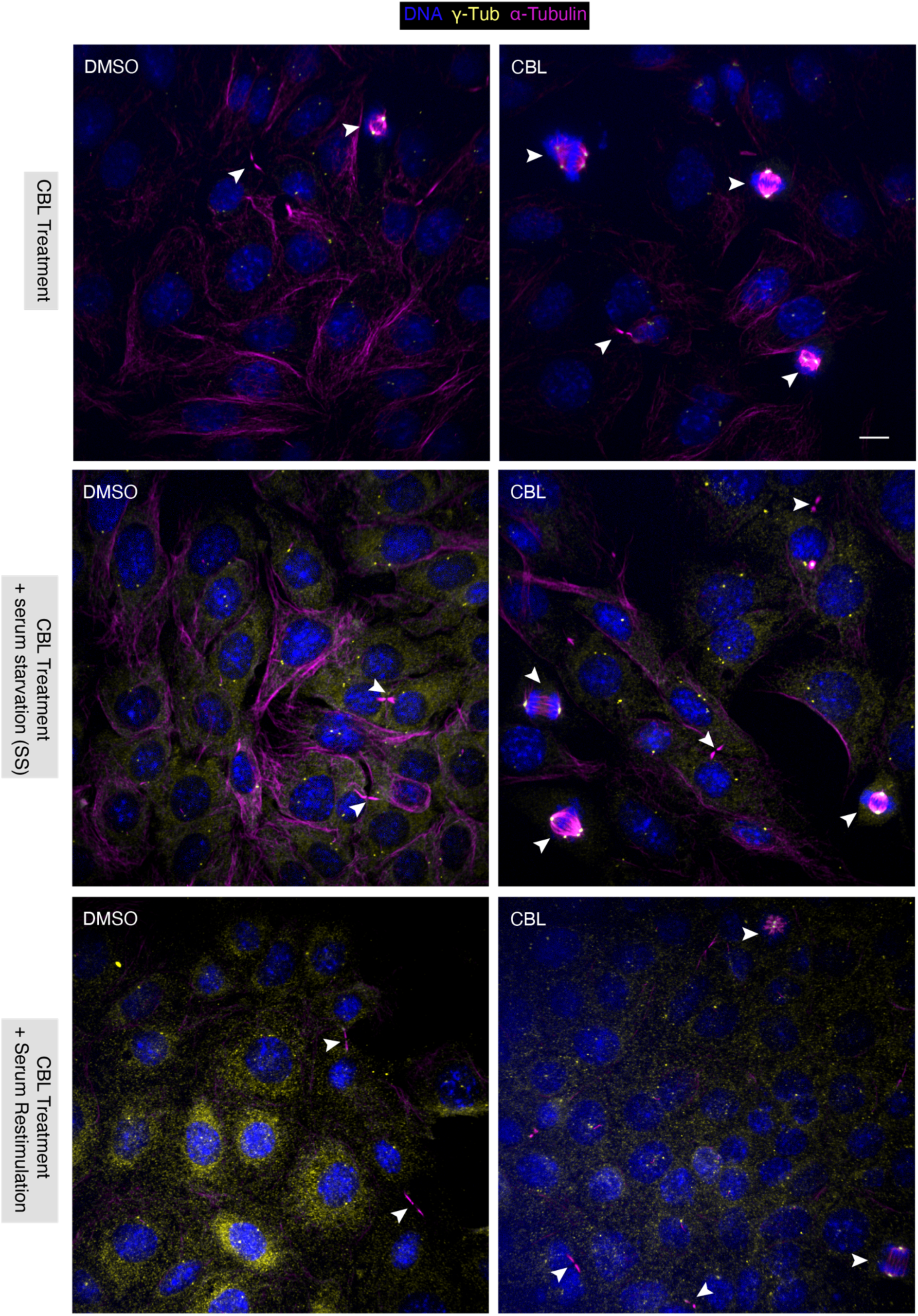
Carbaryl-induced mitotic arrest in IMCD3 cells across treatment conditions. Representative immunofluorescence micrographs corresponding to Figure 3A. White arrowheads indicate mitotic cells and chromatin/anaphase bridges identified within each field. Scale bar: 10 µm.

**Figure S5.**
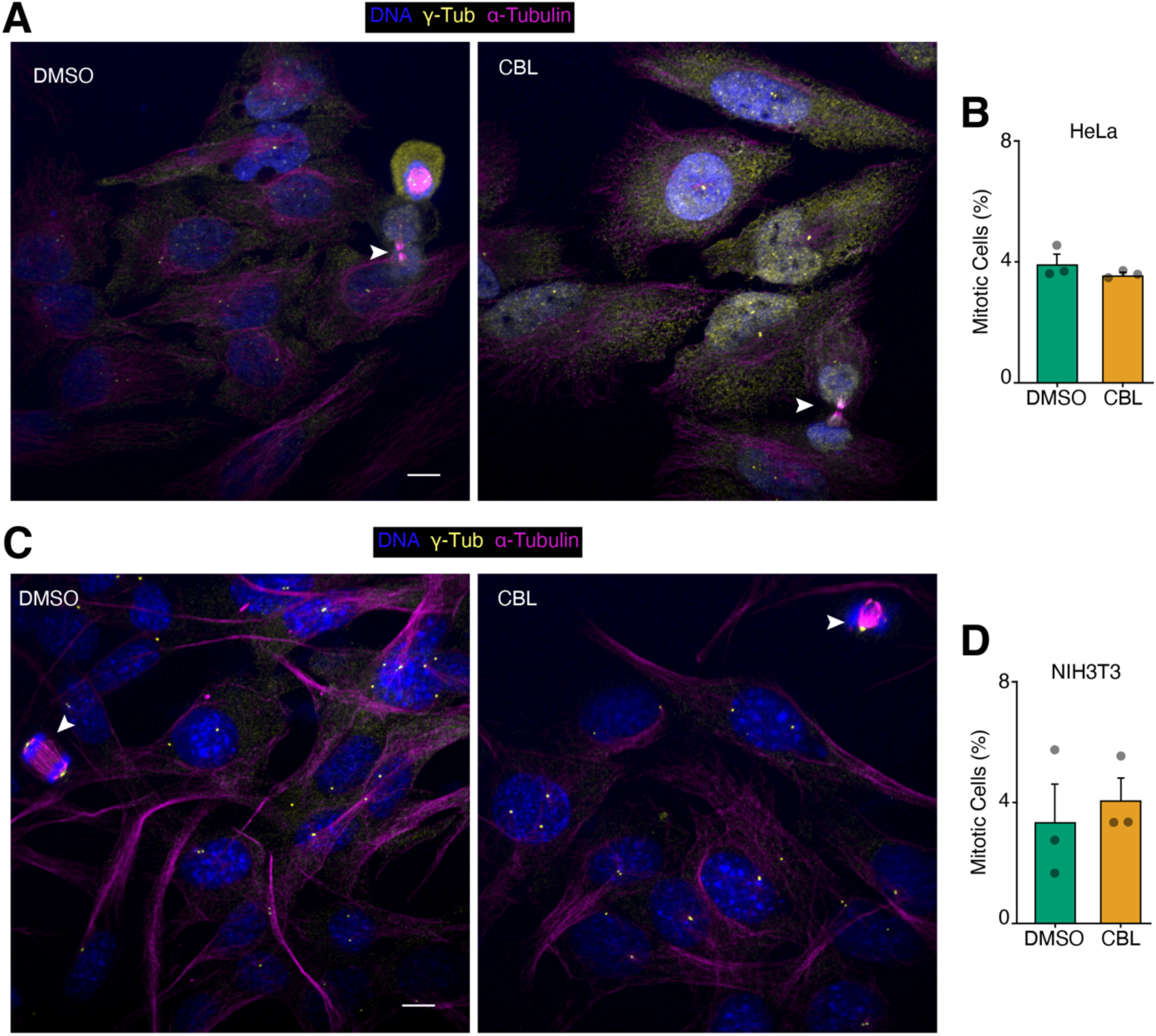
Carbaryl does not induce a mitotic arrest in HeLa or NIH3T3 cells. **(A)** Representative immunofluorescence images of HeLa cells stained for DNA (blue), γ-tubulin (yellow), and α-tubulin (magenta) following DMSO or carbaryl treatment. White arrowheads indicate mitotic cells. Scale bar: 10 µm. **(B)** Quantification of the percentage of mitotic cells in DMSO- versus carbaryl-treated HeLa cells. Data represent the mean ± SEM from three independent experiments. **(C)** Representative immunofluorescence images of NIH3T3 cells stained for DNA (blue), γ-tubulin (yellow), and α-tubulin (magenta) following DMSO or carbaryl treatment. White arrowheads indicate mitotic cells. Scale bar: 10 µm. **(D)** Quantification of the percentage of mitotic cells in DMSO- versus carbaryl-treated NIH3T3 cells. Data represent the mean ± SEM from three independent experiments. Data represent the mean ± SEM from three independent experiments. Asterisks denote statistical significance as determined by unpaired Student’s t-test: **** p ≤ 0.0001; *** p ≤ 0.001; ** p ≤ 0.01; * p ≤ 0.05.

**Figure S6.**
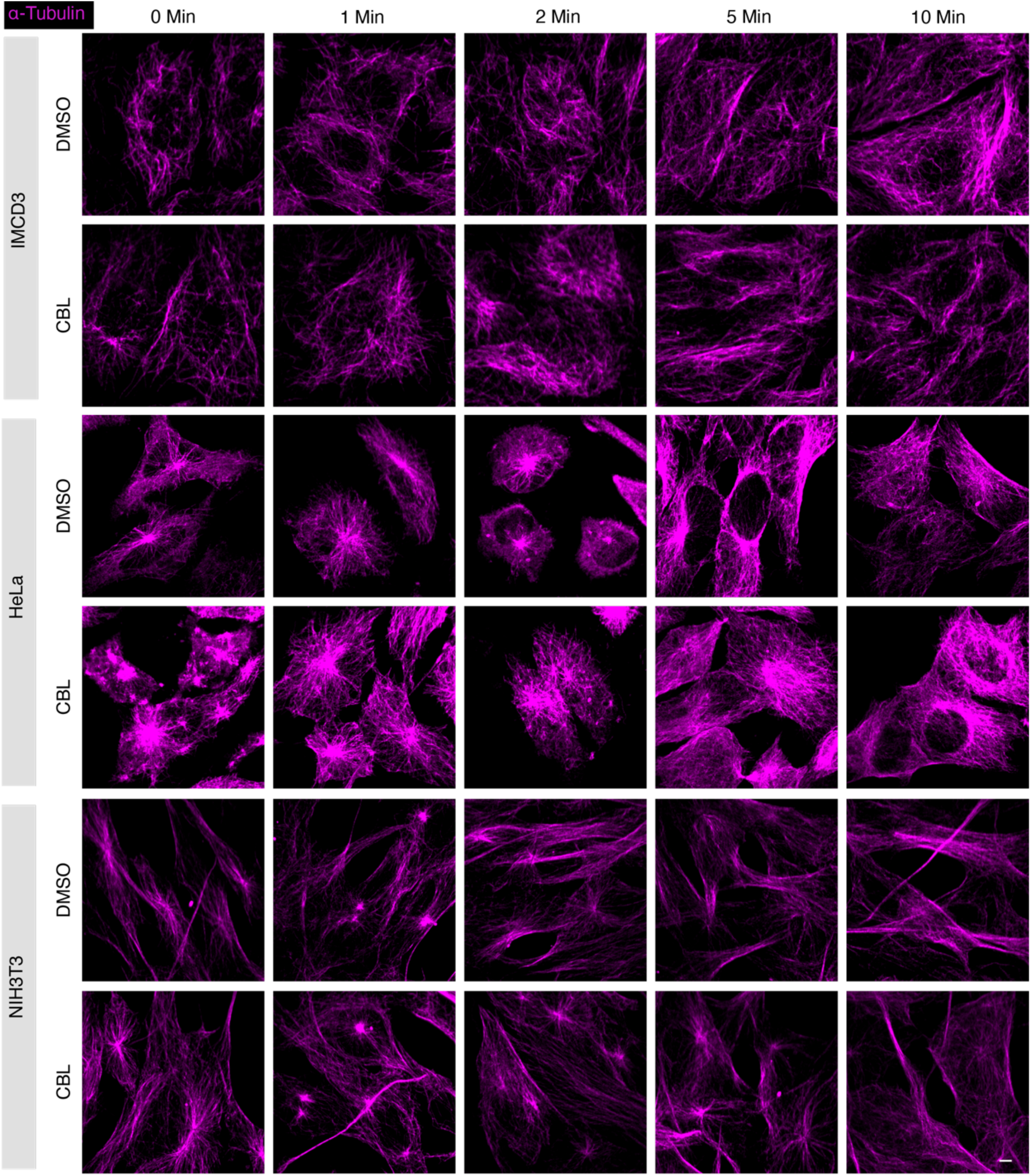
Carbaryl does not alter microtubule regrowth kinetics across IMCD3, HeLa, and NIH3T3 cells. Representative immunofluorescence images of α-tubulin (magenta) in IMCD3 (top), HeLa (middle), and NIH3T3 (bottom) cells treated with DMSO or carbaryl. Microtubules were depolymerized by cold treatment, followed by warming at 37 ℃ to permit regrowth, and cells were fixed at 0, 1, 2, 5, and 10 minutes post-rewarming to assess microtubule network reformation. Across all three cell lines, carbaryl-treated cells show microtubule regrowth kinetics comparable to DMSO controls at each time point, progressing from a depolymerized/aster-like state immediately after cold exposure (0–2 min) to a re-established filamentous microtubule network by 5–10 minutes. Scale bar, 5 µm.

**Figure S7:**
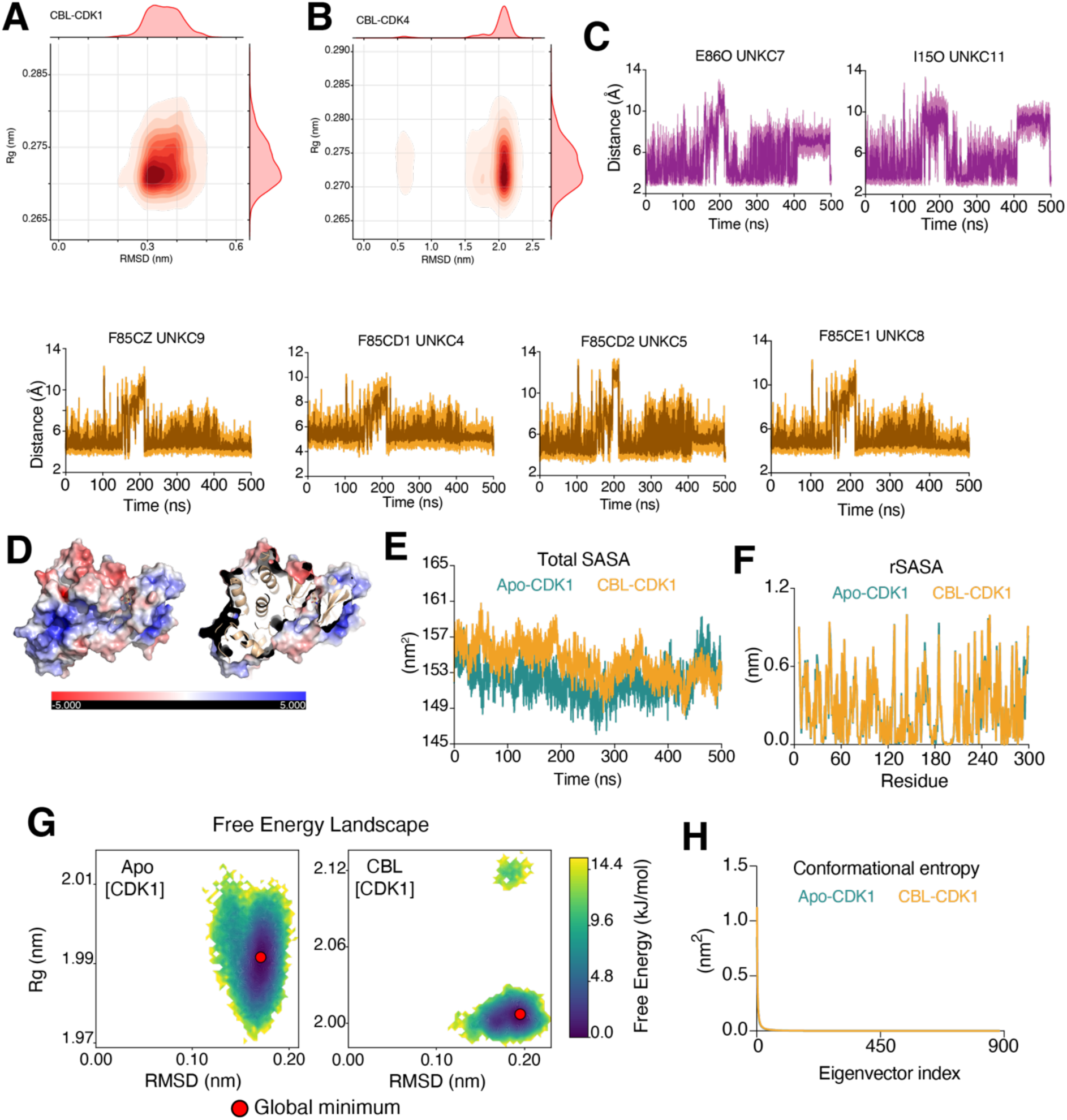
Carbaryl interacts stably with CDK1 throughout the MD simulation. **(A** and **B)** RMSD versus radius of gyration (Rg) plot of carbaryl, demonstrating the structural stability and compactness of carbaryl with CDK1 and CDK4 throughout the 500 ns MD simulation. **(C)** Distance-time plots of key CDK1 residues interacting with carbaryl over the 500 ns simulation. Orange traces represent π-π interactions; purple traces represent hydrophobic interaction distances (Å). The full MD simulation trajectory is provided in Movie **S1**. **(D)** Electrostatic surface potential representation of the CDK1: carbaryl complex. The inset shows a zoomed view of the electrostatic surface potential at the carbaryl binding site. **(E)** Total solvent-accessible surface area (SASA) of apo (green) and carbaryl-bound (wheat) CDK1 over the simulation, reflecting the overall change in solvent accessibility upon carbaryl binding. **(F)** Relative solvent-accessible surface area (rSASA) expressed as a percentage for individual CDK1 residues, illustrating changes in solvent accessibility between the apo and carbaryl-bound forms. **(G)** Two-dimensional free energy landscape (2D-FEL) of CDK1 in the apo and carbaryl-bound states, illustrating the conformational energy states sampled during the simulation. A red dot indicates the global energy minimum of each state. **(H)** Conformational entropy analysis of CDK1 in the apo and carbaryl-bound states, showing that carbaryl binding does not alter global conformational entropy.

**Figure S8:**
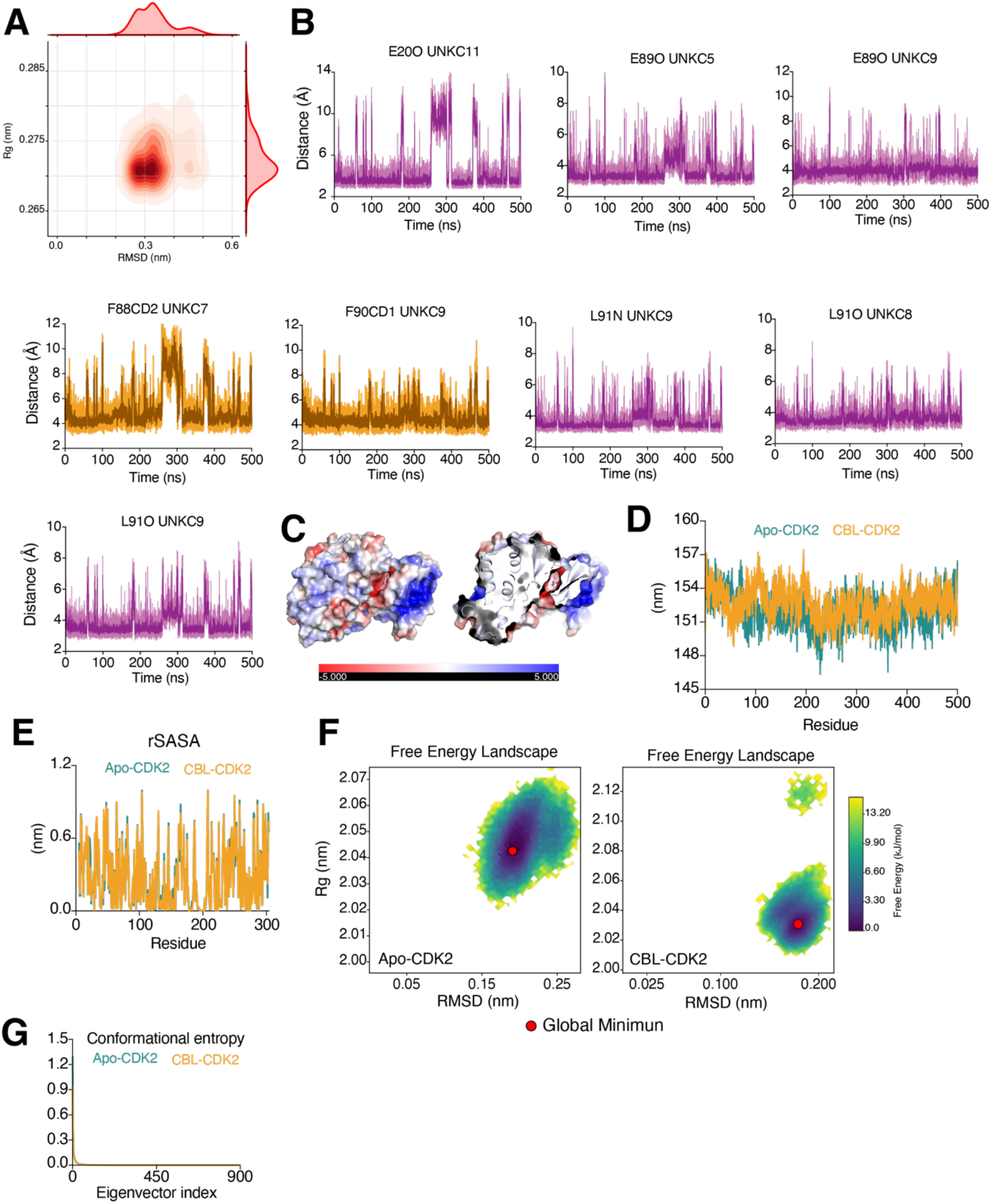
Carbaryl interacts stably with CDK2 throughout the MD simulation. **(A)** RMSD versus radius of gyration (Rg) plot of carbaryl, demonstrating the structural stability and compactness of carbaryl with CDK2 throughout the 500 ns MD simulation. **(B)** Distance-time plots of key CDK2 residues interacting with carbaryl over the 500 ns simulation. Orange traces represent π-π interactions; purple traces represent hydrophobic interaction distances (Å). The full MD simulation trajectory is provided in Movie **S2**. **(C)** Electrostatic surface potential representation of the CDK2: carbaryl complex. The inset shows a zoomed view of the electrostatic surface potential at the carbaryl binding site. **(D)** Total solvent-accessible surface area (SASA) of apo (green) and carbaryl-bound (wheat) CDK2 over the simulation, reflecting the overall change in solvent accessibility upon carbaryl binding. **(E)** Relative solvent-accessible surface area (rSASA) expressed as a percentage for individual CDK2 residues, illustrating changes in solvent accessibility between the apo and carbaryl-bound forms. **(F)** Two-dimensional free energy landscape (2D-FEL) of CDK2 in the apo and carbaryl-bound states, illustrating the conformational energy states sampled during the simulation. A red dot indicates the global energy minimum of each state. **(G)** Conformational entropy analysis of CDK2 in the apo and carbaryl-bound states, showing that carbaryl binding does not alter global conformational entropy.

**Figure S9:**
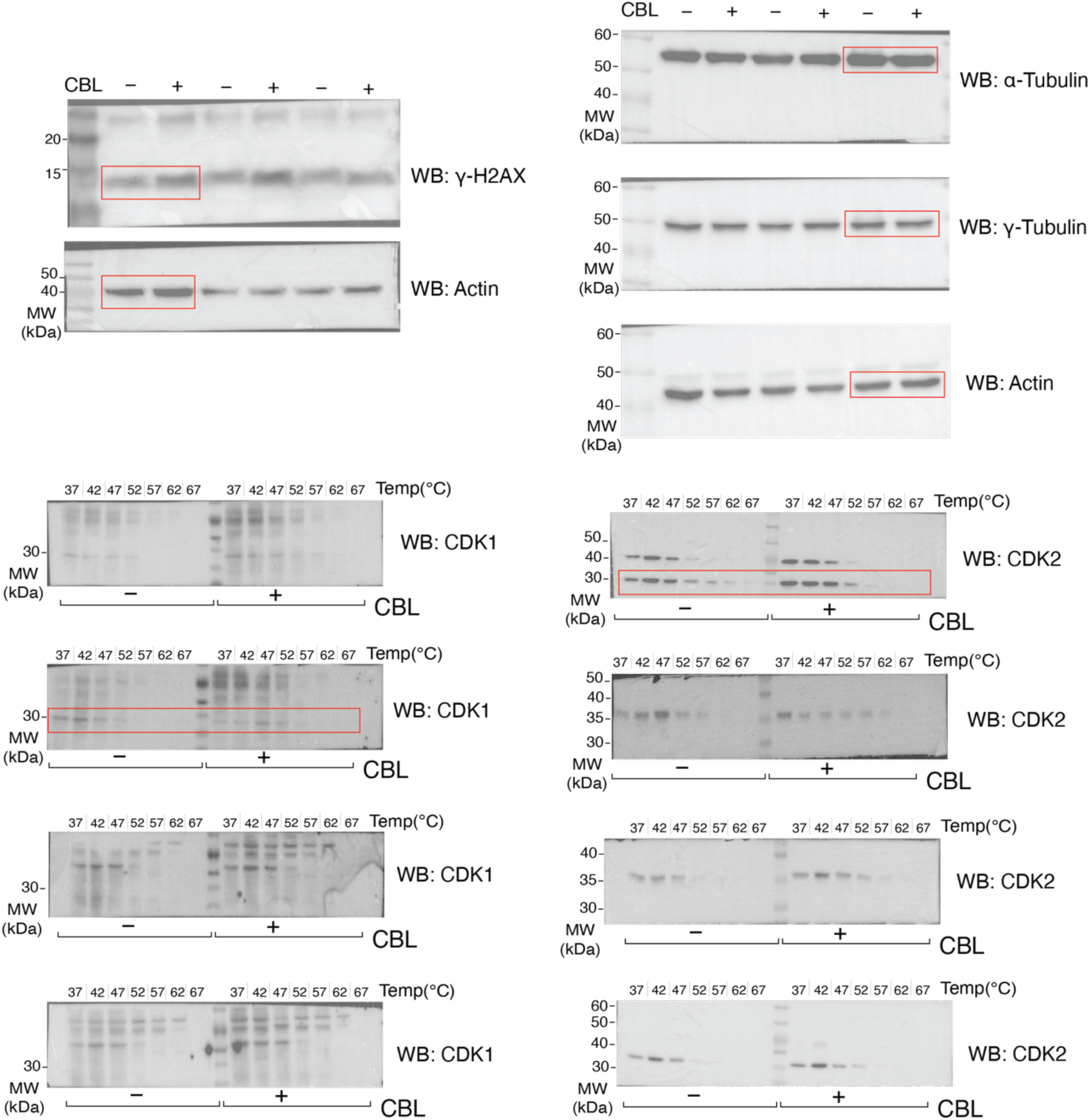
Uncropped Western blots.

**Table S1:** Effect of carbaryl on non-target models.

| Organism/Model | Exposure type | Observed effect | References |
| --- | --- | --- | --- |
| Humans (occupationally exposed workers) | Chronic occupational exposure | Reproductive toxicity, sperm abnormalities, genotoxicity | [34] |
| Human HepG2 cells | *In vitro* | Oxidative stress and DNA damage | [35] |
| Rats | Acute and chronic exposure | Cholinergic toxicity, neurotoxicity, reproductive effects at higher doses | [36] |
| Mice | Chronic exposure | Neurotoxicity and systemic toxicity at high doses | [36] |
| *Caenorhabditis elegans* | Laboratory exposure | Oxidative stress, reduced survival | [37] |
| Zebrafish embryos | Developmental exposure | Teratogenicity, developmental toxicity | [38] |
| Zebrafish larvae | Environmentally relevant concentrations | Behavioral impairment | [39] |
| Fish gill cells (Flounder) | *In vitro* | Cytotoxicity and genotoxicity | [38] |
| Cyanobacteria (*Calothrix brevissima*) | Environmental exposure | Growth inhibition and oxidative stress | [40] |
| Earthworm (*Metaphire posthuma*) | Soil exposure | Reproductive and behavioral toxicity | [41] |

